# CIP2A is required for mitotic recruitment of the SLX1/XPF/MUS81 tri-nuclease complex to replication stress-induced DNA lesions to maintain genome integrity

**DOI:** 10.1101/2025.04.03.647079

**Authors:** Lauren de Haan, Sietse J. Dijt, Alejandro García López, Dan Ruan, Femke J. Bakker, Marieke Everts, Harry Warner, Frank N. Mol, H. Rudolf de Boer, Pim J. Huis in ’t Veld, Bert van de Kooij, Rifka Vlijm, Marcel A.T.M. van Vugt

## Abstract

Perturbed DNA replication can lead to incompletely replicated DNA when cells enter mitosis and can interfere with chromosome segregation. Cells therefore require mechanisms to resolve these lesions during mitosis. The CIP2A-TOPBP1 complex is described to function as a molecular tether that connects fragmented DNA molecules. However, whether CIP2A also functions in processing of incompletely replicated DNA remained unclear. We show that CIP2A-TOPBP1 forms large filamentous structures at sites of incomplete DNA replication during mitosis, which recruit the SMX tri-nuclease complex members SLX4, MUS81 and ERCC1/XPF. These structures form in proximity to sites of mitotic DNA synthesis, although CIP2A is not required for mitotic DNA synthesis. In addition to its globular and coiled-coil domain, the unstructured C-terminal domain of CIP2A is essential for CIP2A-TOPBP1 filamentous structure formation and recruitment of the SMX complex. *BRCA1*^-/-^ and *BRCA2*^-/-^ cells have increased mitotic DNA lesions that recruit CIP2A and SLX4. We show that the C-terminal part of CIP2A is required for survival of *BRCA2*^-/-^ cells. Moreover, SLX4 is crucial for genome stability in *BRCA2*^-/-^ cells. Combined, we demonstrate that CIP2A-TOPBP1 recruits the SMX complex during mitosis, which is required to resolve mitotic DNA lesions, allows faithful chromosome segregation and maintain viability of *BRCA2*^-/-^ cells.

## Introduction

DNA replication is a vital process in all biological systems. To faithfully duplicate the entire genome, DNA replication adheres to a strict temporal and spatial program^1^. Even under physiological conditions, DNA replication is challenged by intrinsic properties of some genomic regions that are difficult to replicate^2^, while also chemical modifications of DNA can perturb replication^3–5^. In cancer cells, however, many additional processes lead to perturbed DNA replication^6,7^. For instance, oncogene activation leads to uncoordinated firing of replication origins, causing collisions between the transcription and replication machinery^8^ and depletion of the nucleotide pool^9,10^. In addition, cancer-associated DNA repair defects also disrupt DNA replication. Specifically, mutations in DNA repair genes *BRCA1*, *BRCA2* and *FANCD2* cause instability of stalled replication forks and incomplete replication^11–14^.

Cells are equipped with several mechanisms to deal with perturbed DNA replication. For instance, when replication forks cannot progress, dormant replication origins in the vicinity of the stalled replication fork are fired to complete replication^15^. In addition, translesion synthesis (TLS) polymerases can replace the replicative polymerases to continue replication^16^. Finally, cell cycle checkpoints prevent mitotic entry when cells have extensive amounts of incompletely replicated DNA or unrepaired DNA lesions^17^. Nevertheless, despite the presence of these protective mechanisms, cancer cells frequently enter mitosis with incompletely replicated DNA^12,18,19^.

After cells have entered mitosis, several mechanisms are available to resolve incompletely replicated DNA. First, DNA replication can be completed during mitosis by ‘mitotic DNA synthesis’ (MiDAS), which involves a dedicated DNA polymerase complex, and resembles RAD52-dependent break-induced replication (BIR)^20,21^. Second, several DNA endonucleases are activated during mitosis (e.g. SLX1, MUS81, XPF)^22^, or get access to chromosomes upon mitotic breakdown of the nuclear membrane (i.e. GEN1)^23^. These endonucleases can separate joint DNA molecules, allowing faithful chromosome segregation and completion of mitosis^24,25^. Third, incompletely replicated DNA fragments that remain unresolved early during mitosis can form ‘ultrafine DNA bridges’ (UFBs) during anaphase^26,27^. UFBs arise at sites of incomplete DNA replication, and are marked by the recruitment of the PICH DNA translocase^28^. In turn, PICH recruits the BLM-TopoIII-RMI1/2 complex^26,27^ and RIF1^29^. UFBs originating from incompletely replicated DNA are converted into single-stranded DNA bridges^29,30^, which likely facilitates UFB breakage and allows chromosome segregation at the cost of genome instability^30^. The molecular regulation of these various mechanisms remains largely elusive, and it is unclear whether these pathways act in parallel or consecutively, and whether these mechanisms are part of an integrated response to incompletely replicated DNA.

The CIP2A-TOPBP1 complex was recently demonstrated to be recruited to mitotic DNA lesions^31,32^. The CIP2A-TOPBP1 complex tethers the two ends of double-stranded DNA breaks (DSBs) and prevents mis-segregation of acentric chromosome fragments^31–34^. Importantly, CIP2A is essential for survival of *BRCA1/2* mutant cells^31^, which accumulate mitotic DNA lesions^12,19^. Interestingly, whereas TOPBP1 can be recruited to DNA lesions both during interphase and mitosis^35–38^, CIP2A is sequestered in the cytoplasm during interphase, and is only recruited to DSBs upon nuclear envelop breakdown during mitosis^31,32^. Recruitment of CIP2A and TOPBP1 to mitotic DSBs depends on the γH2AX adaptor protein MDC1^39^, however DNA lesions induced by the replication polymerase inhibitor aphidicolin (APH) appear to be MDC1-independent^31,32^, showing that multiple mechanisms exist for CIP2A-TOPBP1 recruitment. APH treatment induces mitotic DNA lesions, including joint DNA molecules that likely require additional processing when compared to mitotic DSBs. However, it is currently unclear if the CIP2A-TOPBP1 complex has functions beyond the tethering of DNA ends, and functions in processing of mitotic DNA lesions induced by perturbed replication.

In this study, we used stimulated emission Depletion (STED) super-resolution microscopy and structure-function analysis of CIP2A to study mitotic CIP2A-TOPBP1 structure formation. We find that the CIP2A-TOPBP1 complex mediates recruitment of the SMX tri-nuclease complex to sites of replication-associated DNA lesions during mitosis. Recruitment of the SMX complex by CIP2A-TOPBP1 allows for processing of DNA lesions induced by perturbed replication, and is essential for genome stability in *BRCA2*^-/-^ cells.

## Results

### DNA lesions induced by perturbed replication and irradiation result in similar CIP2A-TOPBP1 structures during mitosis

Analysis of the cellular response to mitotic DNA damage is frequently studied using irradiation of mitotic cells to induce DSBs. Since transfer of late-stage replication intermediates into mitosis is a cancer cell-intrinsic source of mitotic DNA damage, we compared CIP2A recruitment to replication stress-induced and irradiation-induced mitotic DNA damage. To this end, hTERT immortalized human retinal pigmented epithelial RPE1 *TP53*^-/-^ cells were treated with the polymerase inhibitor aphidicolin (APH) or with irradiation (IR). Analysis of mitotic foci formation of CIP2A in response to low-dose APH (200 nM) or low-dose IR (0.25 Gy) confirmed that the majority of CIP2A foci co-localizes with γH2AX in mitosis, and that CIP2A recruitment to mitotic DNA lesions occurs independently of the source of DNA damage (Fig. 1A, B and Suppl. Fig. 1A)^31,32^. To examine the effects of CIP2A inactivation on the response to mitotic DNA damage, RPE1 *TP53*^-/-^ *CIP2A*^-/-^ clones were established using CRISPR-Cas9 (Suppl. Fig. 1B). CIP2A loss prevented TOPBP1 localization at DNA selectively during mitosis, which verifies the lack of functional CIP2A expression, and allowed us to selectively study the role of the mitotic CIP2A-TOPBP1 complex (Suppl. Fig. 1B-D)^31,32^. In line with previous studies, *CIP2A*^-/-^ cells showed elevated levels of micronuclei, already in untreated conditions^31–33^. Importantly, the amount of micronucleated *CIP2A*^-/-^ cells strongly increased upon APH and IR treatment (Fig. 1C, Suppl. Fig. 1E), showing that CIP2A prevents missegregation of chromosome fragments into micronuclei in response to various sources of mitotic DNA damage.

**Figure 1:**
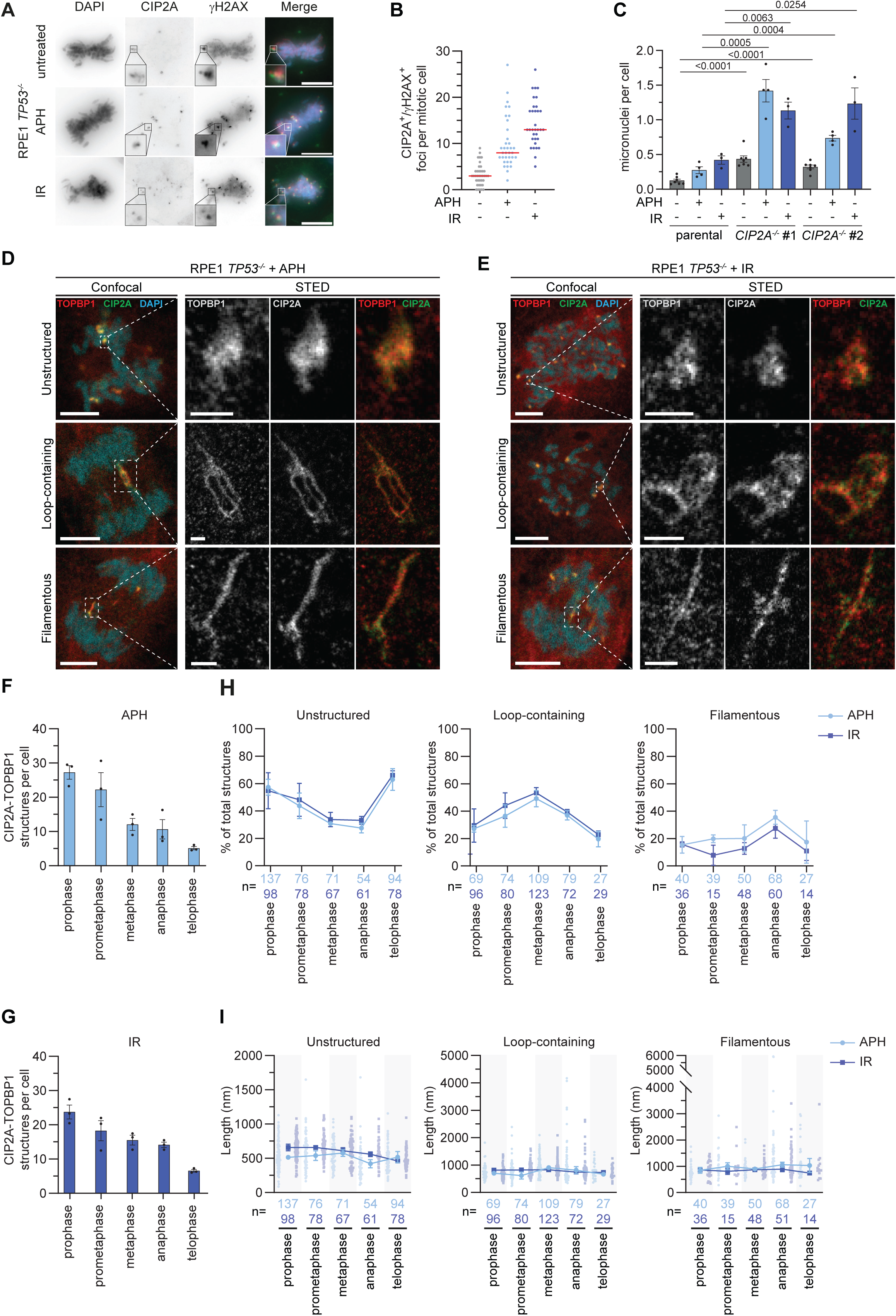
DNA damage-induced mitotic CIP2A-TOPBP1 structures. **(A)** RPE1 *TP53*^-/-^ cells were either left untreated or treated with aphidicolin (APH, 200 nM, 20 h) or ionizing radiation (IR, 0.25 Gy). Cells were stained for DAPI (blue), CIP2A (green) and γH2AX (red). Scale bar represents 10 µm. **(B)** Quantification of co-localizing CIP2A and γH2AX foci per mitotic cell for cells as treated as described in A. Individual values and medians of n>30 cells per condition are shown. **(C)** RPE1 *TP53^-/-^* cells and *CIP2A^-/-^* clones were either left untreated or treated with APH (200 nM, 24 h) or IR (3 Gy) and analyzed 72 h later. Micronuclei per cell were quantified. The bars represent the mean and standard error of the mean (SEM) from three biologically independent experiments with n≥64 cells per experimental condition. P-values were calculated using two-tailed unpaired t-test. **(D, E)** RPE1 *TP53^-/-^* cells were treated with APH (200 nM, 20 h, panel D) or IR (0.25 Gy, panel E). Representative images of three classes of mitotic structures are shown. Left panels show the confocal overview images, and the right three panels show STED images of a single CIP2A-TOPBP1 complex. Cells were stained for DAPI (blue), CIP2A (green) and TOPBP1 (red). Scale bar represents 5 µm (confocal) or 500 nm (STED). STED images are Wiener deconvolved. Raw data is shown in Suppl. Fig 1G, H. **(F)** Quantification of CIP2A-TOPBP1 structures per cell per mitotic phase for cells treated as described in panel D. Number of cells quantified per mitotic phase from three biologically independent experiments: prophase (n=9), prometaphase (n=9), metaphase (n=19), anaphase (n=18), telophase (n=29). The numbers of CIP2A-TOPBP1 structures per telophase may be an overestimation of the actual number, as only cells with CIP2A-TOPBP1 structures were measured. Bars represent mean and SEM of three experiments. **(G)** Quantification of CIP2A-TOPBP1 structures per cell per mitotic phase for cells treated with IR as described in panel E. Number of cells quantified per mitotic phase from three biologically independent experiments: prophase (n=11), prometaphase (n=10), metaphase (n=14), anaphase (n=13), telophase (n=19). Bars represent mean and SEM of three biologically independent experiments. **(H)** Quantification of unstructured, loop-containing and filamentous structures per mitotic phase for indicated treatments, as observed with STED microscopy for cells treated as described in from panels D and E. Percentages compared to the total structures are indicated. Mean, SEM and n, which reflects the total number of observed structures per phase, from three biologically independent experiments are shown per mitotic phase and per treatment. **(I)** Quantification of the size of unstructured, loop-containing and filamentous structures per mitotic phase for indicated treatments, as observed with STED microscopy for cells treated as described in panels D and E. Structure size was determined by measuring the length of the longest axis through the entire structure as shown in Suppl. Fig. 1J. Individual values, the average of medians per experiment, along with SEM, are plotted per mitotic phase, ‘n’ represents the total number of observed structures per phase from three biologically independent experiments.

We subsequently explored the role of the ATM and ATR DNA damage response (DDR) kinases in CIP2A recruitment. ATR activity was not required for APH-induced induction of mitotic CIP2A foci, nor was it required for CIP2A foci formation in untreated cells (Suppl. Fig. 1F). Rather, ATR inhibition elevated the number of CIP2A foci in untreated cells, suggesting that ATR inhibition leads to increased transmission of DNA lesions into mitosis (Suppl. Fig. 1F). ATM inhibition did not influence the number of CIP2A foci in either untreated or APH-treated cells. However, ATM inhibition reduced the number of IR-induced CIP2A foci (Suppl. Fig. 1F). These results show that CIP2A marks different types of lesions, with differential upstream DNA damage signaling requirements.

To obtain a more detailed view of mitotic CIP2A-TOPBP1 structures, we employed stimulated emission depletion (STED) microscopy. This type of super-resolution imaging offers a spatial resolution of 30-50 nm, compared to confocal microscopy with a resolution of around 250 nm^40,41^. STED microscopy revealed three distinct classes of CIP2A-TOPBP1 structures: unstructured complexes, loop-containing structures and filamentous structures (Fig. 1D, E, Suppl. Fig. 1G, H). These three classes of structures occurred both in response to APH or IR treatment, which all showed a high degree of CIP2A and TOPBP1 co-localization within these structures, indicating that these proteins are less than 30 nm apart from each other (Fig. 1D, E, Suppl. Fig. 1G-I). We observed a clear reduction in the number of CIP2A-TOPBP1 structures during mitotic progression (Fig. 1F, G). No obvious differences in prevalence of these structures between APH or IR treatment was observed. Instead, the prevalence of the distinct structures depended on the mitotic stage (Fig. 1H). Whereas the majority of the CIP2A-TOPBP1 complexes in prophase and prometaphase showed an unstructured morphology (45-55%), in metaphase the loop-containing structures were more prevalent (51%) (Fig. 1H). While the filaments only constituted a constant minor fraction of the total amount of structures (15%), during anaphase their prevalence strongly increased (31%). In telophase, the remaining CIP2A-TOPBP1 complexes mostly showed an unstructured morphology (Fig. 1H). Examination of the structure size by measuring the length of the longest axis of the entire structure (Suppl. Fig. 1J) showed that the median size of unstructured forms ranged from 505-656 nm in prophase to 453-472 nm in telophase (Fig. 1I), with IR-induced complexes being slightly larger than APH-induced complexes. The median size of loop-containing structures ranged from 827-903 nm, with no obvious difference between APH or IR-induced structures. Filamentous structures showed a median size range between 730-870 nm in prophase to 724-825 nm in telophase, with no obvious difference between IR and APH conditions (Fig. 1I). Of note, whereas the median size of loop-containing and filamentous structures was similar between APH- and IR-treated cells and between the different mitotic stages, the size range of these structures became larger in later mitotic stages (e.g. range of IR-induced filamentous structures in anaphase: 389-3388 nm). Combined, these analyses illustrate the emergence of large CIP2A-TOPBP1 structures, that show similar morphologies despite being formed in response to different sources of DNA damage (Fig. 1I).

### CIP2A interacts with the SMX complex upon perturbed replication

Although APH and IR treatment both resulted in micronuclei formation in *CIP2A*^-/-^ cells and induced the formation of similar CIP2A-TOPBP1 structures, the underlying DNA lesions differ and likely require differential processing during mitosis. To explore if the CIP2A-TOPBP1 complex has differential interacting partners upon APH versus IR treatment, endogenous CIP2A was immunoprecipitated in parental RPE1 *TP53*^-/-^ or *CIP2A*^-/-^ cells and associated proteins were analyzed. The established CIP2A interactors TOPBP1^31,32^ and MDC1^32^ were identified by western blot analysis, validating the approach (Suppl. Fig. 2A). Mass spectrometry analysis of CIP2A immunoprecipitations identified several additional interacting partners (Fig. 2A, B, Suppl. Fig. 2B, C. We identified the XPF endonuclease (also called ERCC4) specifically in APH treatment conditions, (Fig. 2A, B; Suppl. Fig. 2B, C; Suppl. Table 1). In line with the mass spectrometry analysis, XPF co-localized with CIP2A at APH--induced foci (Fig. 2C). In line with our mass spec results, in untreated and IR-treated conditions most CIP2A foci were XPF-negative, whereas >50% of APH-induced CIP2A foci were XPF-positive (Fig. 2C, Suppl. Fig. 2D). These findings imply that complex formation of CIP2A-TOPBP1 with XPF depends on the type of DNA lesion. Importantly, CIP2A was required to recruit XPF to APH-induced DNA lesions, as XPF foci were absent in *CIP2A*^-/-^ cells (Fig. 2D, H). Based on these findings, we further focused on APH-induced mitotic DNA lesions.

**Figure 2:**
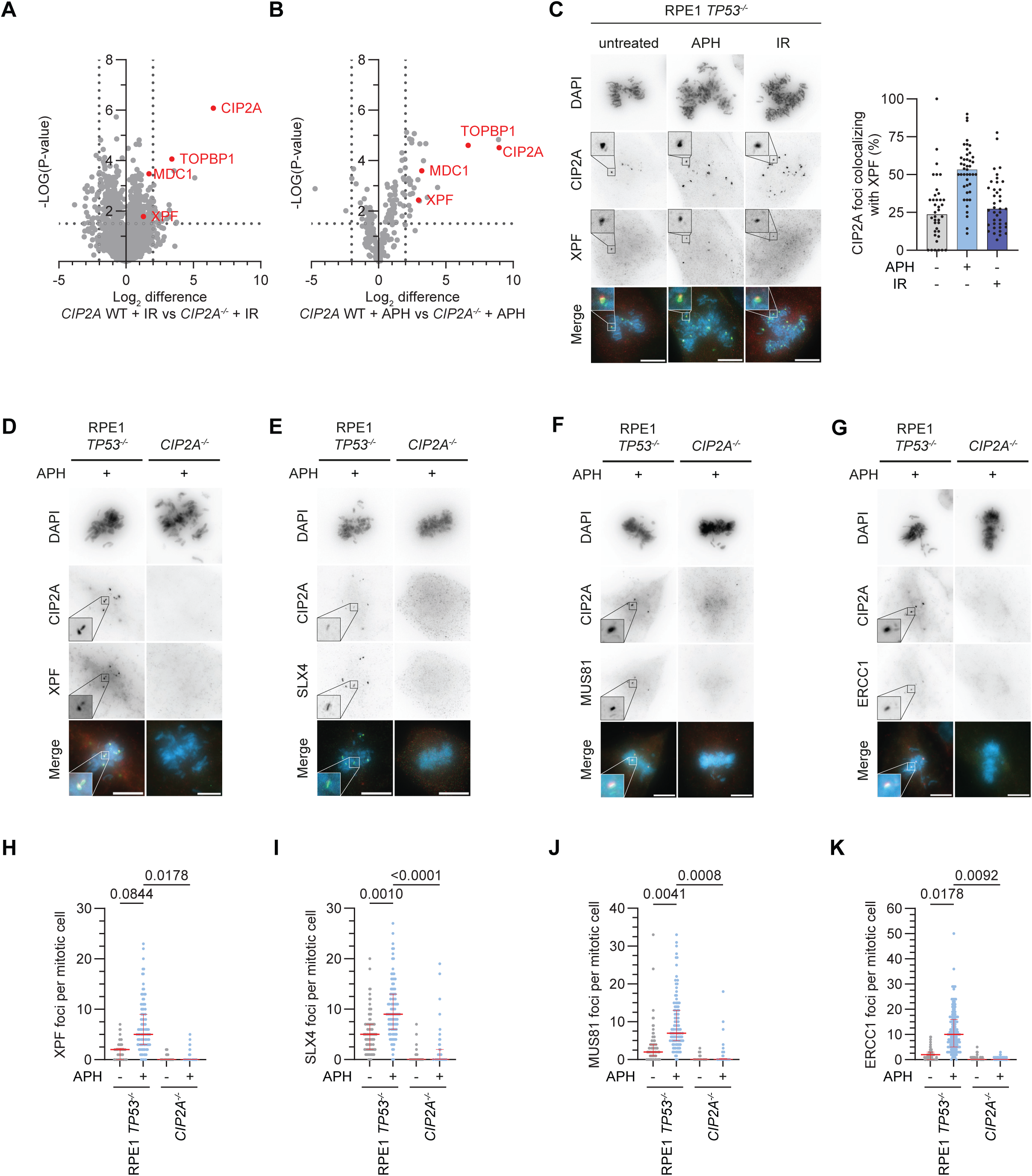
CIP2A is required for mitotic recruitment of the SLX4-ERCC1-XPF-MUS81 complex. **(A, B)** Mass spectrometry analysis of mitotic CIP2A interactors in RPE1 *TP53^-/-^* parental versus *CIP2A^-/-^* cl#1 cells upon treatment with IR (panel A, 5 Gy) or with APH (panel B, 200 nM, 20 h). Proteins highlighted in red are enriched after IR or APH treatment. **(C)** Left panel: representative images of RPE1 *TP53^-/-^* cells stained for DAPI (blue), CIP2A (green) and XPF (red) in either untreated conditions, or upon treatment with APH (200 nM, 20 h) or IR (0,25 Gy). Scale bar represents 10 µm. Right panel: quantification of the percentage of CIP2A foci per mitotic cell that co-localize with XPF. Bar represents the median of one experiment with n>35 cells per experimental condition. **(D)** RPE1 *TP53^-/-^* and *CIP2A^-/-^* cl#1 cells were treated with APH (200 nM, 20 h). Representative images of treated cells stained for DAPI (blue), CIP2A (green) and XPF (red). Scale bar represents 10 µm. **(E)** Representative images of RPE1 *TP53^-/-^* cells and *CIP2A^-/-^*cl#1 cells treated with APH (200 nM, 20 h). Cells were stained for DAPI (blue), CIP2A (red) and SLX4 (green). Scale bar represents 10 µm. **(F)** RPE1 *TP53^-/-^* cells and *CIP2A^-/-^* cl#1 cells were left untreated or treated with APH (200 nM, 20 h). Representative images of treated cells stained for DAPI (blue), CIP2A (red) and MUS81 (green) in untreated condition or after treatment with APH (200 nM, 20 h). Scale bar represents 10 µm. **(G)** RPE1 *TP53^-/-^* cells and *CIP2A^-/-^*cl#1 cells were left untreated or treated with APH (200 nM, 20 h). Representative images of treated cells stained for DAPI (blue), CIP2A (red) and ERCC1 (green), in untreated conditions or after treatment with APH (200 nM, 20 h). Scale bar represents 10 µm. **(H)** Quantification of mitotic XPF foci in RPE1 *TP53^-/-^* cells and *CIP2A^-/-^* cl#1 cells either left untreated or treated as described in D. Individual values, medians and interquartile range of three experiments with n>30 cells per experimental condition are plotted. P-values were calculated using two-way ANOVA with Šidák’s multiple comparisons test on median values per experiment. **(I)** Quantification of mitotic SLX4 foci in RPE1 *TP53^-/-^* cells and *CIP2A^-/-^* cl#1 cells either untreated or treated as described in panel E. Individual values, medians and interquartile range of three experiments with n>30 cells per experimental condition are plotted. P-values were calculated using two-way ANOVA with Šidák’s multiple comparisons test on the medians per experiment. **(J)** Quantification of mitotic MUS81 foci in RPE1 *TP53^-/-^* cells and *CIP2A^-/-^* cl#1 cells either left untreated or treated as described in panel F. Individual values, medians and interquartile range of three experiments with n≥27 cells per experiment are plotted. P-values were calculated using two-way ANOVA with Šidák’s multiple comparisons test on the medians per experiment. **(K)** Quantification of mitotic ERCC1 foci in RPE1 *TP53^-/-^* cells and *CIP2A^-/-^* cl#1 cells either left untreated or treated as described in panel G. Individual values, medians and interquartile range of three experiments with n≥28 cells per experimental condition are plotted. P-values were calculated using two-way ANOVA with Šidák’s multiple comparisons test on the medians per experiment.

During mitosis, the structure-specific XPF nuclease along with its co-factor ERCC1 is part of the SMX tri-nuclease complex, which further consists of the SLX4 scaffold and the other structure-specific nuclease SLX1 and MUS81-EME1^42^. The SMX complex has been described to resolve joint DNA molecules, including stalled replication forks and HR intermediates such as Holliday junctions, to enable proper chromosomes segregation in anaphase^24,42–45^. Immunofluorescence microscopy analysis showed that other partners of the SMX complex (i.e. SLX4, MUS81 and ERCC1) also co-localized with CIP2A (Fig. 2E-G,). Importantly, like XPF, mitotic foci formation of SLX4, MUS81 and ERCC1 foci was significantly increased upon APH treatment, and largely absent in *CIP2A*^-/-^ cells (Fig. 2E-G, I-K), demonstrating that the CIP2A-TOPBP1 complex is required for recruitment of the SMX complex during mitosis. To confirm that CIP2A recruits the SMX complex rather than its components individually, SLX4 was depleted in APH-treated cells. SLX4 depletion did not affect CIP2A foci formation (Suppl. Fig. 2E, F, H), whereas SLX4 depletion completely prevented the recruitment of MUS81 to CIP2A foci (Suppl. Fig. 2F, G, I, J). Combined, these data show that CIP2A-TOPBP1 acts upstream of and is required for SMX complex recruitment in mitosis.

### CIP2A-TOPBP1 forms a mitotic scaffold for the SMX complex at sites of perturbed DNA replication

To investigate how the SMX complex is locally positioned at mitotic CIP2A-TOPBP1 structures, STED microscopy analysis of SLX4, ERCC1 and MUS81 was performed. SLX4, ERCC1 and MUS81 showed a punctate localization pattern onto CIP2A structures, with locally enriched recruitment (Fig. 3A-C. Suppl. Fig. 3A-C). Of note, the SMX complex members SLX4, ERCC1 and MUS81 all locate at CIP2A structures (ranging between 77%-97% of CIP2A structures), independently of CIP2A organization (unstructured forms, loop-containing or filamentous structures) at APH-induced DNA lesions (Fig. 3D-F). Line profile analysis showed that SMX complex members localize along the entire CIP2A structure, with some SMX complex members occasionally found at regions with lower CIP2A intensity (Suppl. Fig. 4A). Combined, these data suggest that the mitotic CIP2A-TOPBP1 structures form a scaffold that recruits and localizes the SMX complex to sites of DNA lesions.

**Figure 3:**
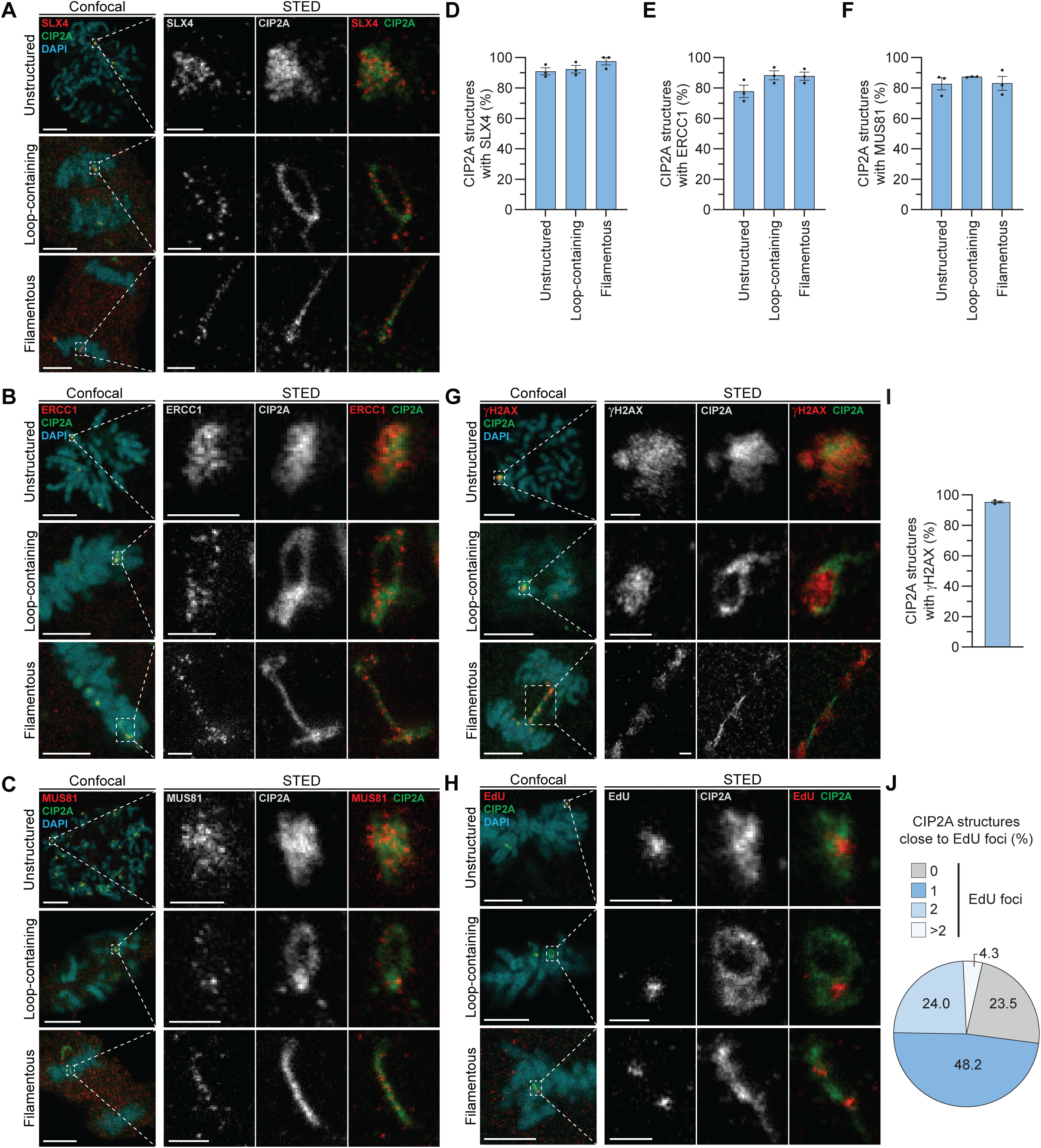
Localization of SMX nucleases at mitotic CIP2A structures. **(A-C)** RPE1 *TP53^-/-^* cells were treated with APH (200 nM, 20 h) and stained for DAPI (blue), CIP2A (green) and SLX4 (red, panel A), ERCC1 (red, panel B), MUS81 (red, panel C). A representative image of each of the CIP2A structure organizations (unstructured, loop-containing, filamentous) are shown. Scale bar represents 5 µm (confocal) or 500 nm (STED). STED images are Wiener deconvolved. Raw data is shown in Suppl. Fig. 3A-C. **(D-F)** Quantification of the percentage of mitotic CIP2A structures that co-localize with SLX4 (panel D), ERCC1 (panel E) or MUS81 (panel F) as observed with STED microscopy for each of the CIP2A structure organizations (unstructured, loop-containing, filamentous) for cells treated as described in panels A-C. Bars represent means and SEM of three experiments with n>75 structures per experiment. **(G)** RPE1 *TP53^-/-^* cells were treated with APH (200 nM, 20 h) and stained for DAPI (blue), CIP2A (green) and γH2AX (red). Scale bar represents 5 µm (confocal) or 500 nm (STED). STED images are Wiener deconvolved. Raw data is shown in Suppl. Fig. 3D. **(H)** RPE1 *TP53^-/-^* cells were treated with APH (200 nM, 20 h), pulsed with EdU during mitotic entry and stained for DAPI (blue), CIP2A (green) and EdU (red). Scale bar represents 5 µm (confocal) or 500 nm (STED). STED images are Wiener deconvolved. Raw data is shown in Suppl. Fig 3E. **(I)** Quantification of mitotic CIP2A structures that are positive for γH2AX as observed by STED microscopy for cells treated as described in panel G. Bars represent means and SEM for three experiments with n>75 structures per experiment. **(J)** Quantification of mitotic CIP2A structures that are either negative (0 foci) or positive for 1, 2 or more than 2 EdU foci as observed by STED microscopy for cells treated as described in panel I. Pie chart represents the means of three experiments with n>75 structures per experiment.

We next analyzed whether CIP2A structures formed at sites of ongoing DNA replication at mitotic entry. To this end, cells were pulsed with the synthetic nucleotide analogue EdU during mitotic entry, and incorporated EdU or γH2AX was visualized along with CIP2A. For unstructured, and filamentous CIP2A structures, γH2AX located at the CIP2A structures, with γH2AX appearing as a cloud surrounding CIP2A structures (Fig. 3G, Suppl. Fig. 3D, 4B). Conversely, loop-containing CIP2A structures predominantly showed γH2AX staining enclosed inside the CIP2A loops (Fig. 3G, Suppl. Fig. 3D, 4B). EdU foci were mainly observed within CIP2A loop structures, and localized to those parts of the CIP2A structures that showed lower CIP2A intensity (Fig. 3H, Suppl. Fig. 3E, 4C). The large majority of CIP2A structures were positive for both γH2AX and EdU (Fig. 3I, J). Of note, filamentous CIP2A structures occasionally bridged two EdU foci (Fig. 3H, Suppl. Fig. 3E). For approximately half of APH-induced CIP2A structures (48.2%), we identified one EdU focus in close proximity, whereas a smaller percentage (24.0%) of CIP2A structures assembled in the vicinity of two EdU foci (Fig. 3J). Together, these data reveal intricate structures of a CIP2A-containing macromolecular scaffold that recruits the SMX complex, and forms at sites of DNA damage in the vicinity of mitotic DNA synthesis.

### CIP2A is not required for mitotic DNA synthesis

Previous studies showed mitotic DNA synthesis at sites of under-replicated DNA in a process called MiDAS^21^. As CIP2A localizes to sites of EdU incorporation, we investigated if CIP2A was required for MiDAS. Analysis of mitotic EdU foci in parental RPE1 *TP53*^-/-^ cells, *CIP2A*^-/-^ cells, and *CIP2A*^-/-^ cells reconstituted with full length V5-tagged CIP2A (Suppl. Fig. 5A) showed that CIP2A is not required for MiDAS in RPE1 *TP53*^-/-^ cells (Fig. 4A, B). Previously, short-term depletion of MUS81, SLX4 or EME1 in U2OS cells was reported to impair MiDAS^21^, however studies in RPE1 *TP53*^-/-^ cells showed that MUS81 was not required for MiDAS induced by APH^46^. In line with our finding that CIP2A is not essential for MiDAS, we confirmed that depletion of MUS81 did not affect APH-induced MiDAS in RPE1 *TP53*^-/-^ cells (Suppl. Fig. 5B). To extend these findings to other cell line models, CIP2A was depleted in a panel of triple-negative breast (TNBC) cell lines. Specifically, *CIP2A* was edited using CRISPR/Cas9-in MDA-MB-231 and HCC38 cells (Fig. 4C-E, Suppl. Fig. 5C, D), or depleted by shRNA in BT549 cells (Fig. 4E, Suppl. Fig. 5E). *CIP2A* inactivation prevented mitotic SLX4 foci formation (Fig. 4C, D). In contrast, in all cell lines tested, CIP2A depletion did not prevent EdU foci formation in response to APH treatment (Fig. 4C-E). Combined, these data show that CIP2A is not essential for MiDAS.

**Figure 4:**
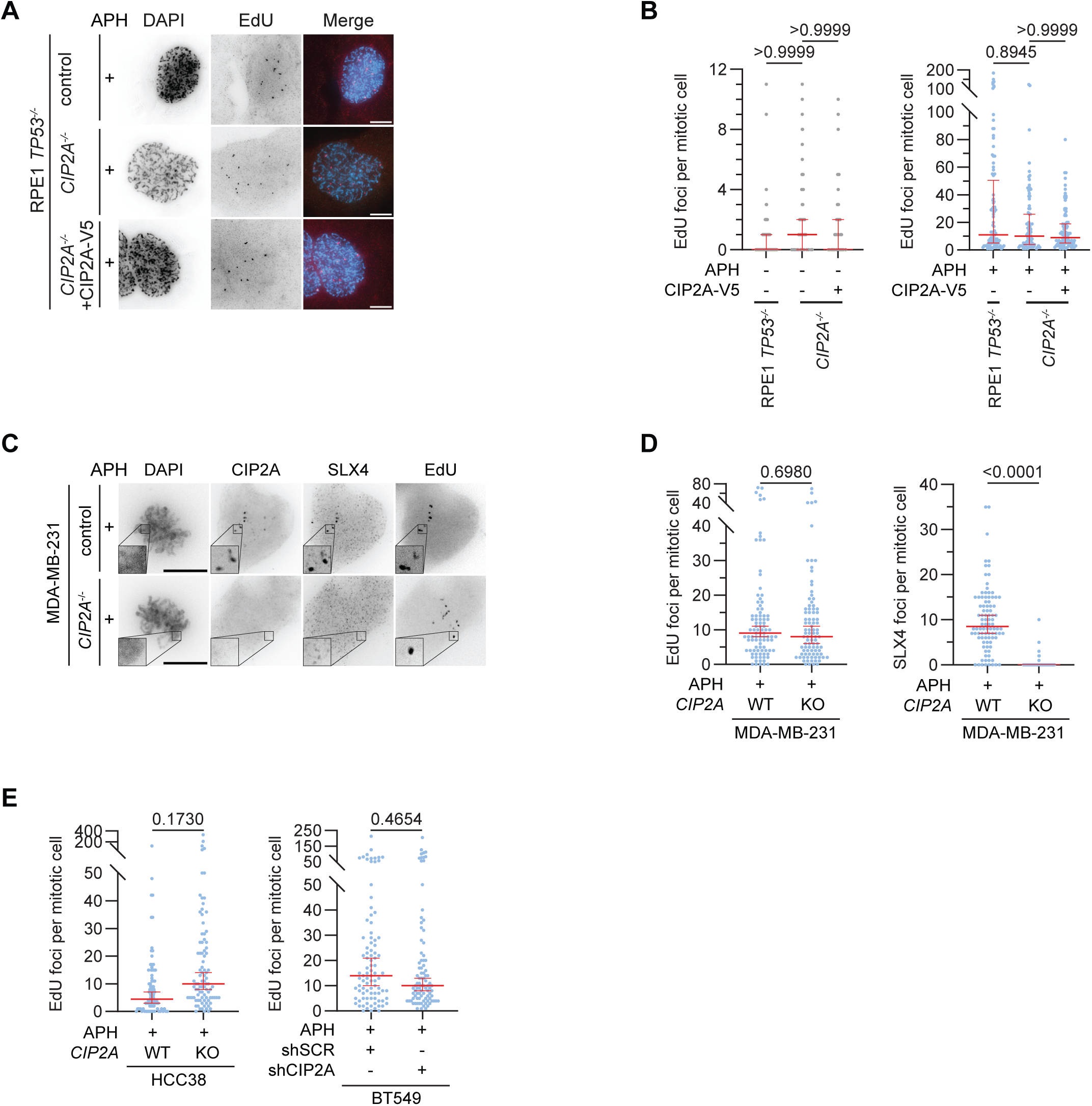
CIP2A is not essential for mitotic DNA synthesis. **(A)** RPE1 *TP53^-/-^*, *CIP2A^-/-^* cl#1 cells and CIP2A-V5 reconstituted *CIP2A^-/-^*cl#1 cells were treated with APH (200 nM, 20 h), pulsed with EdU during mitotic entry and stained for DAPI (blue) and EdU (red) Scare bar represents 10 µm. **(B)** Quantification of EdU foci per mitotic cell in untreated RPE1 *TP53^-/-^* cells, *CIP2A^-/-^* cl#1 cells, and *CIP2A^-/-^* cl#1 cells reconstituted with full CIP2A WT-V5 or cells treated as described in panel A. Individual values, medians and interquartile range of three experiments with n>28 cells per experimental condition are plotted. P-values were calculated using ordinary one-way ANOVA with Šidák’s multiple comparisons test on the medians per experiment. **(C)** Wildtype or *CIP2A^-/-^* MDA-MB-231 cells were treated with APH (200 nM, 20 h) and pulsed with EdU during mitotic entry. Representative images of MDA-MB231 cells stained for DAPI, CIP2A, SLX4, EdU are shown. Scale bar represents 10 µm. **(D)** Quantification of EdU and SLX4 foci per mitotic cell for cells treated as described in panel C. Individual values, medians and interquartile range of three experiments with n≥29 cells per experimental condition are plotted. P-values were calculated using two-tailed unpaired t-test on the medians per experiment. **(E)** Wildtype or *CIP2A^-/-^*HCC38 cells, and doxycycline-treated BT549 cells with doxycycline-inducible scramble (shSCR) or CIP2A (shCIP2A) shRNA were treated with APH (200 nM, 20 h), and pulsed with EdU during mitotic entry. Quantification of EdU foci per mitotic cell in HCC38 and BT549. Individual values, medians and interquartile range of three experiments with n>30 cells per experimental conditions are plotted. P-values are calculated using two-tailed unpaired t-test on the medians per experiment.

### The C-terminal domain of CIP2A is required for the formation of mitotic CIP2A-TOPBP1 structures and SMX complex recruitment

The CIP2A protein structure analysis showed three distinct domains: an N-terminal globular domain (aa 1-617), an alpha-helical coiled-coil domain (aa 618-876), and a conserved C-terminal unstructured domain (aa 877-905). AlphaFold predicted CIP2A to form a stable homodimer, with a coiled-coil shaft with an uncertain - and possibly flexible - position of the N-terminal globular domain onto the coiled-coil shaft (Fig. 5A, B, Suppl. Fig. 6A, B). AlphaFold predictions further suggested that a CIP2A homodimer binds to two TOPBP1 proteins (Fig. 5A, B, Suppl. Fig. 6A). This CIP2A-TOPBP1 interaction involves residues 755-860 of TOPBP1, which likely interact with both the globular domain and coiled-coil of CIP2A (Fig. 5A, B, Suppl. Fig. 6B).

**Figure 5:**
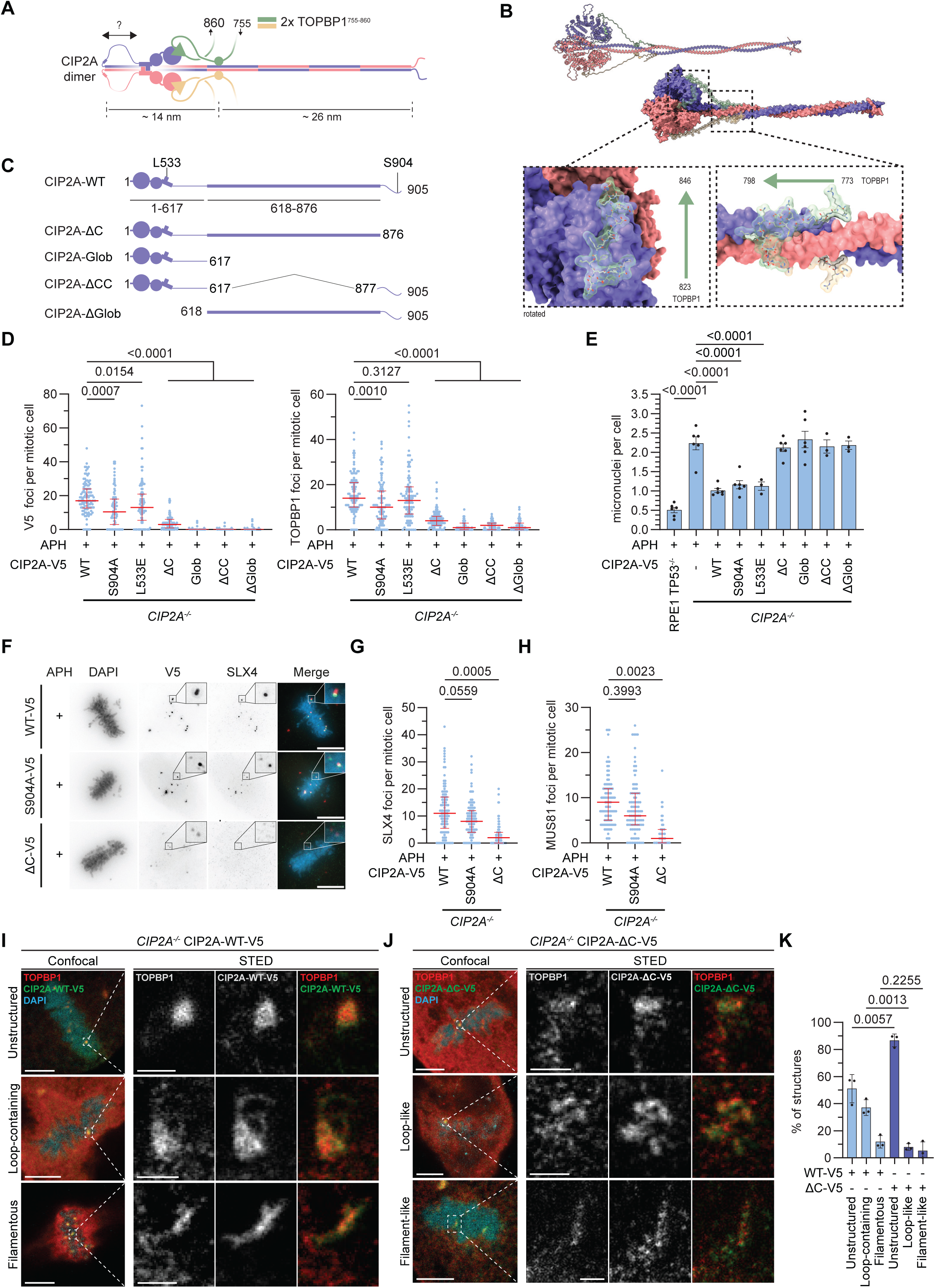
Structure-function analysis of CIP2A. **(A)** CIP2A is predicted to form a stable homodimer, involving a globular domain with a long coiled-coil. Two predicted binding sites of TOPBP1 are indicated, along with an uncertain, and possibly flexible, position of the N-terminal regions onto the coiled-coil shaft. **(B)** A cartoon and surface representation of the predicted CIP2A:CIP2A dimer structure with two copies of TOPBP1 755-860, and a magnification of predicted TOPBP1:CIP2A binding interfaces. **(C)** Schematic representation of full length (1-905) CIP2A (WT), the reported dimerization site (L533), the PLK1 phosphorylation site (S904), the globular domain (1-617), the alpha-helical coiled-coil (618-876), and the unstructured C-terminal domain (877-905). Schematic representation of V5-tagged CIP2A variants: CIP2A lacking the unstructured C-terminal domain (‘CIP2A-ΔC’), CIP2A with only globular domain (‘CIP2A-Glob’), CIP2A lacking the alpha-helical coiled-coil (‘CIP2A-ΔCC’) and CIP2A lacking the globular domain (‘CIP2A-ΔGlob’). **(D)** Quantification of V5 foci and TOPBP1 foci per mitotic cell in RPE1 *TP53^-/-^ CIP2A^-/-^* cl#1 cells reconstituted with indicated CIP2A-V5 cDNAs, treated with APH (200 nM, 20 h). Individual values, medians and interquartile range of three experiments with n>30 cells per experimental condition are plotted. P-values were calculated using ordinary one-way ANOVA with Šidák’s multiple comparisons test on the medians per experiment. **(E)** Quantification of micronuclei per cell in parental RPE1 *TP53^-/-^* cells, *CIP2A^-/-^*cl#1 cells and *CIP2A^-/-^* cl#1 cells reconstituted with indicated CIP2A cDNAs, treated with APH (200 nM, 20 h). Individual values, medians and interquartile range of at least three experiments with n≥85 cells per experimental condition are plotted. P-values were calculated using ordinary one-way ANOVA with Dunnett’s multiple comparison test. **(F)** *CIP2A^-/-^* cells reconstituted with CIP2A-WT, CIP2A-S904A or CIP2A-ΔC were treated with APH (200 nM, 20 h), and stained for DAPI (blue), V5 (red) and SLX4 (green). Scale bar represents 10 µm. **(G)** Quantification of SLX4 foci per mitotic cell in *CIP2A^-/-^* cl#1 reconstituted with indicated mutants for cells treated as described in panel F. Individual values, medians and interquartile range of three experiments with n>30 cells per experimental condition are plotted. P-values were calculated using ordinary one-way ANOVA with Šidák’s multiple comparisons test on the medians per experiment. **(H)** Quantification of MUS81 foci per mitotic cell in *CIP2A^-/-^* cl#1 reconstituted with either CIP2A-WT, CIP2A-S904A or CIP2A-ΔC after treatment with APH (200 nM, 20 h). Individual values, medians and interquartile range of three experiments with n>30 cells per experimental condition are plotted. P-values were calculated using ordinary one-way ANOVA with Šidák’s multiple comparisons test on the medians per experiment. **(I)** Representative STED microscopy images of DAPI (blue), V5 (green) and TOPBP1 (red) structures in *CIP2A^-/-^* cells reconstituted with CIP2A-WT for the three classes of structures (unstructured, loop-containing, and filamentous) after APH treatment (200 nM, 20 h). Scale bar represents 5 µm (confocal) or 500 nm (STED). STED images are Wiener deconvolved. Raw data is shown in Suppl. Fig. 7D **(J)** Representative STED microscopy images of DAPI (blue), V5 (green) and TOPBP1 (red) structures in *CIP2A^-/-^* cells reconstituted with CIP2A-ΔC for the three classes of structures (unstructured, loop-containing, and filamentous) after APH treatment (200 nM, 20 h). Scale bar represents 5 µm (confocal) or 500 nm (STED). STED images are Wiener deconvolved. Raw data is shown in Suppl. Fig. 7E. **(K)** Quantification of unstructured, loop-containing or loop-like, and filamentous or filament-like structures in *CIP2A^-/-^*cells reconstituted with either CIP2A-WT or CIP2A-ΔC. Bars represent the mean and SEM of three experiments with n>15 structures per experimental condition. P-values were calculated using two-tailed unpaired t-test.

To investigate which domains of CIP2A are required for complex formation of CIP2A-TOPBP1 and for SMX recruitment, a panel of CIP2A deletion mutants was analyzed. *CIP2A*^-/-^ cells were reconstituted with either full length CIP2A (WT: CIP2A 1-905-V5), a deletion mutant lacking the C-terminal unstructured region (ΔC: CIP2A 1-876-V5), a mutant only having the globular domain (Glob: CIP2A 1-617-V5), a mutant lacking the coiled-coil domain (ΔCC: CIP2A Δ618-878-V5), and a mutant lacking the globular domain (ΔGlob: CIP2A 618-905-V5) (Fig. 5C). Additionally, we included the CIP2A L533E mutant that was previously reported to interfere with CIP2A dimerization^47^, and a S904A mutant to disrupt a previously reported putative PLK1 phosphorylation site (Suppl. Fig. 6C)^48^. All CIP2A mutants showed stable expression, except for CIP2A ΔGlob (Suppl. Fig. 5A). Immunofluorescence microscopy analysis demonstrated that CIP2A-WT, CIP2A-S904A, and CIP2A-L533E formed clear mitotic foci which co-localized with TOPBP1. In line with our AlphaFold predictions that TOPBP1 interacts with both the globular and coiled-coil domain of CIP2A, both mutants lacking the coiled-coil or the globular domain precluded CIP2A-TOPBP1 foci formation (Fig. 5D). Strikingly, cells expressing CIP2A-ΔC only formed very faint CIP2A-TOPBP1 foci, when compared to cells expressing CIP2A-WT (Suppl. Fig. 7A, B).

To explore the functional consequences of these CIP2A mutants, micronuclei formation after APH treatment was investigated. *CIP2A* deletion increased micronuclei formation in RPE1 cells, which was rescued upon expression of CIP2A-WT, S904A and L533E, whereas expression of CIP2A-ΔC, or the mutants that did not form any CIP2A-TOPBP1 foci, did not rescue this phenotype (Fig. 5E). We next tested whether the reduced CIP2A-TOPBP1 foci formation of CIP2A-ΔC expressing cells coincided with defective SLX4 and MUS81 recruitment. In these analyses, CIP2A-S904A was included as S904 resides in the 29 aa C-terminal region that is lacking in CIP2A-ΔC. Whereas SLX4 and MUS81 foci formation was only modestly and non-significantly decreased in cells expressing CIP2A-S904A, it was strongly reduced in cells expressing CIP2A-ΔC (Fig. 5F-H, Suppl. Fig. 7C). Combined, these observations show that the unstructured C-terminal tail of CIP2A is required for the formation of higher order CIP2A-TOPBP1 structures, for the prevention of micronuclei formation and for SMX complex recruitment.

STED microscopy was subsequently used to analyze the role of the C-terminal tail of CIP2A in mitotic CIP2A-TOPBP1 structure formation. Expression of CIP2A-ΔC in *CIP2A^-/-^*cells showed the formation of low intensity CIP2A structures that co-localized with TOPBP1 (Fig. 5I, J, Suppl. Fig. 7D, E). However, formation of loop-containing and filamentous CIP2A-ΔC structures was rare, and the loop-containing and filamentous structures that did form in CIP2A-ΔC expressing cells appeared less ordered than CIP2A-WT structures (Fig. 5I, J, K, Suppl. Fig. 7D, E). Taken together, these data indicate that the C-terminal tail of CIP2A is required for correct organization of mitotic CIP2A-TOPBP1 structures and subsequently SMX recruitment.

### Mitotic CIP2A-SLX4 foci formation in BRCA1 and BRCA2 mutant cells

*BRCA1* and *BRCA2* mutant cells were shown to have increased CIP2A foci in mitosis and loss of BRCA1/2 is synthetic lethal with CIP2A inactivation^31^. This prompted us to investigate mitotic CIP2A foci in isogenic RPE1 and DLD1 cell line panels with WT or mutant *BRCA1/2*. Both *BRCA1*^-/-^ and *BRCA2*^-/-^ cells showed elevated levels of CIP2A foci (Fig. 6A-D). Importantly, a large fraction of CIP2A foci in *BRCA1*^-/-^ and *BRCA2*^-/-^ cells showed co-localization with SLX4 (Fig. 6A-D). Specifically, >65% of CIP2A foci in *BRCA2*^-/-^ cells was positive for SLX4 (Fig. 6D), whereas approximately 30% of the CIP2A foci in *BRCA1*^-/-^ cells was positive for SLX4 (Fig. 6C). Taken together, these data indicate that a large fraction of mitotic CIP2A foci recruit SLX4 in *BRCA1*^-/-^ and *BRCA2*^-/-^ cells, and suggest that mitotic SMX recruitment could be required for survival of these cells.

**Figure 6:**
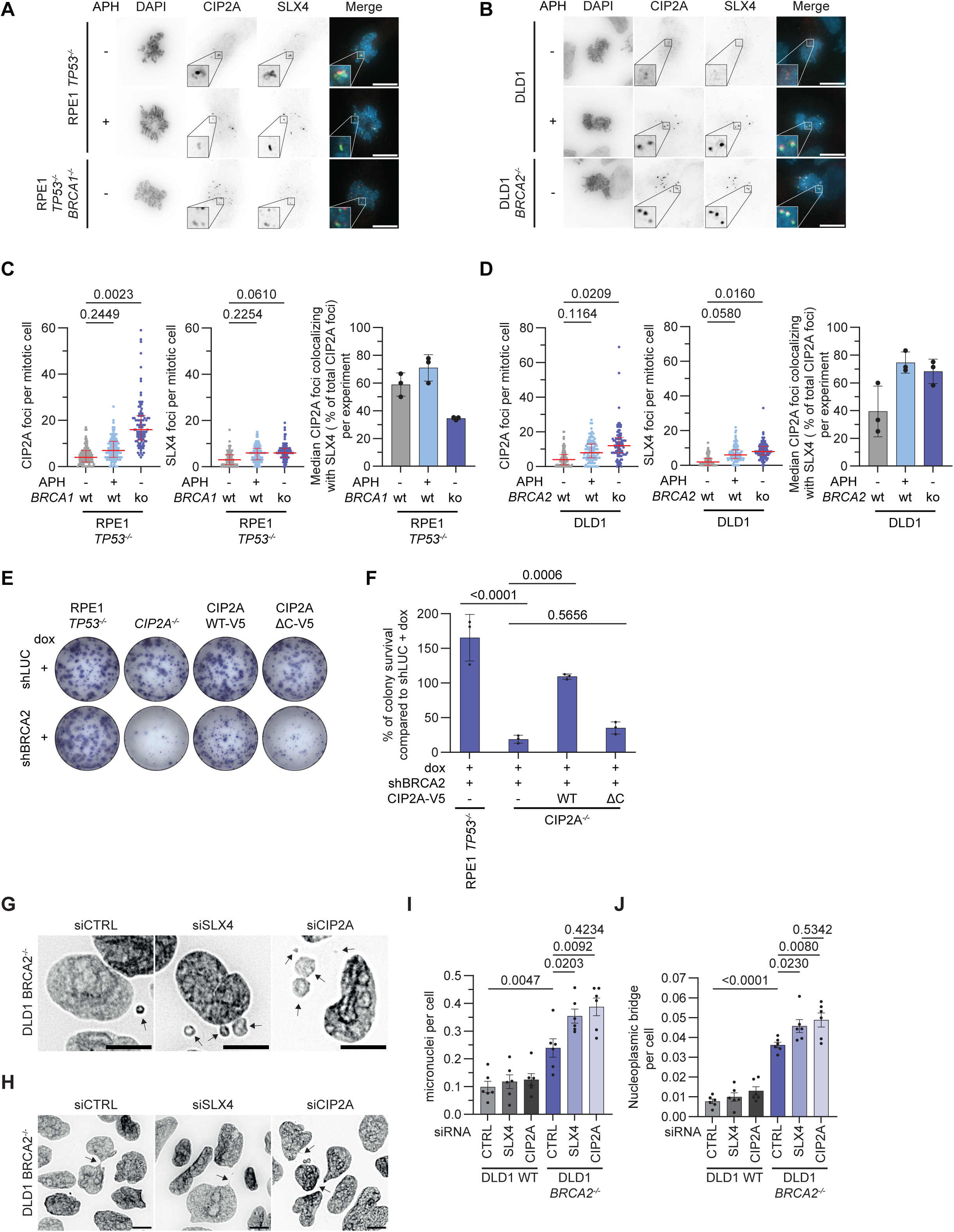
mitotic CIP2A-SLX4 foci in *BRCA1/2* mutant cells. **(A)** Representative images of RPE1 *TP53^-/-^ PAC^-/-^* cells and RPE1 *TP53^-/-^ PAC^-/-^ BRCA1^-/-^* cells, either left untreated or treated with APH (200 nM, 20 h) and stained for DAPI (blue), CIP2A (red) and SLX4 (green). Scale bar represents 10 µm. **(B)** Representative images of DLD1 WT cells and DLD1 *BRCA2^-/-^* cells, either left untreated or treated with APH (200 nM, 20 h), and stained for DAPI (blue), CIP2A (red) and SLX4 (green) in. Scale bar represents 10 µm. **(C)** Quantification of CIP2A and SLX4 foci per mitotic cell for cells treated as described in panel A. Individual values, medians and interquartile range of three experiments with n>30 cells per experimental condition are plotted. P-values were calculated using two-way ANOVA with Šidák’s multiple comparisons test on the medians per experiment. Quantification of CIP2A foci co-localizing with SLX4 for cells treated as described in panel A. Median values per experiment are plotted. Bars represent the mean and SD of three experiments with n>30 cells per experimental condition. **(D)** Quantification of CIP2A and SLX4 foci per mitotic cell for cells treated as described in panel B. Individual values, medians and interquartile range of three experiments with n>30 cells per experimental condition are plotted. P-values were calculated using two-way ANOVA with Šidák’s multiple comparisons test on the medians per experiment. Quantification of the total number of CIP2A foci that co-localize with SLX4 for cells treated as described in panel B. Median values per experiment are plotted. Bars represent the mean and SD of three experiments with n>30 cells per experimental condition. **(E)** Representative images of clonogenic survival assays of doxycycline-treated parental RPE1 *TP53^-/-^* and *CIP2A^-/-^* cl#1 and *CIP2A^-/-^* cl#1 cells, reconstituted with CIP2A-WT or CIP2A-ΔC with doxycycline-inducible luciferase (shLUC) or BRCA2 (shBRCA2) shRNA. **(F)** Quantification of colony survival for cells treated as described in panel E, normalized to doxycycline-treated shLUC-expressing cell lines. Bars represent mean survival and SD of three experiments. P-values were calculated using ordinary one-way ANOVA with Dunnett’s multiple comparison test. **(G-H)** Representative images of DLD1 *BRCA2^-/-^* cells transfected with indicated siRNAs and stained for DAPI. Scale bar represents 10 µm. Arrows indicate either micronuclei (panel G), or nucleoplasmic bridges (panel H). **(I)** Quantification of number of micronuclei per cell for DLD1 WT cells or DLD1 *BRCA2^-/-^* cells, transfected with indicated siRNAs. Bars represent the mean with SEM of 6 experiments with at least n>280 cells per experimental condition. P-values were calculated using two-tailed unpaired t-test. **(J)** Quantification of number of nucleoplasmic bridges per cell for DLD1 WT cells or DLD1 *BRCA2^-/-^* cells transfected with indicated siRNAs. Bars represent the mean with SEM of 6 experiments with at least n>280 cells per experimental condition. P-values were calculated using two-tailed unpaired t-test.

As *BRCA1*^-/-^ and *BRCA2*^-/-^ cells showed an increased number of mitotic CIP2A foci which were positive for SLX4, we investigated if expression of CIP2A-ΔC, which precluded SMX recruitment, is synthetic lethal with BRCA2 loss. To this end, we depleted BRCA2 using doxycycline-inducible shRNAs in parental RPE1 *TP53^-/-^* cells*, CIP2A^-/-^* cells, or *CIP2A^-/-^* cells reconstituted with CIP2A-WT or CIP2A-ΔC (Suppl. Fig. 8A). In line with a previous report^31^, inactivation of *CIP2A* resulted in a strong inhibition of clonogenic outgrowth of BRCA2-depleted RPE1 *TP53^-/-^* cells (Fig. 6E, F). Importantly, viability of BRCA2 depleted cells was rescue by expression of full-length CIP2A, but not by expression of CIP2A-ΔC (Fig. 6E, F). Clearly, formation of higher order CIP2A-TOPBP1 complexes that can recruit the SMX complex is required to resolve the mitotic DNA lesions in *BRCA1/2* mutant cells.

### SLX4 loss phenocopies CIP2A loss in BRCA2-deficient cells

Next, we aimed to investigate whether mitotic recruitment of the SMX complex, like CIP2A, is required for genome stability of BRCA2-deficient cells. To assess this, we depleted either SLX4 or CIP2A in BRCA2-proficient and BRCA2-deficient cells (Suppl. Fig. 8B) and quantified micronuclei formation as a marker of mitotic defects. A significant and similar increase in micronuclei formation was observed in BRCA2-deficient cells upon depletion of either SLX4 or CIP2A, whereas no significant changes were observed in BRCA2-proficient cells (Fig. 6G, I). These findings indicate that SLX4 prevents mitotic errors in BRCA2-deficient cells, to a similar extent as CIP2A.

We further hypothesized that depletion of SLX4 or CIP2A in BRCA2-deficient cells disrupts the processing of under-replicated DNA lesions, resulting in persistent DNA connections following the completion of mitosis. To test this, we quantified the number of nucleoplasmic bridges in BRCA2-proficient and -deficient cells upon SLX4 or CIP2A depletion. Consistent with a role for CIP2A and SLX4 in processing joint molecules that result from perturbed replication, we observed a significant increase in nucleoplasmic bridges upon loss of SLX4 or CIP2A in BRCA2-deficient cells, but not in BRCA2-proficient cells (Fig. 6H, J). Combined, our results show that SLX4, which is recruited during mitosis by CIP2A-TOPBP1, is crucial for preventing excessive genome deterioration in BRCA2-deficient cells.

## Discussion

In this study we describe the formation of CIP2A-TOPBP1 structures in response to DNA damage during mitosis. Importantly, we find that the CIP2A-TOPBP1 scaffold recruits the SMX tri-nuclease complex to sites of under-replicated DNA. Formation of mitotic CIP2A-TOPBP1 structures requires a highly conserved short C-terminal domain of CIP2A, which is also required for SMX recruitment and viability of *BRCA2* mutant cancer cells. Combined, these results demonstrate that the CIP2A-TOPBP1 complex functions beyond DNA damage tethering and is required for mitotic DNA lesion processing.

We observed that both perturbed DNA replication and irradiation lead to CIP2A-TOPBP1 structures with similar morphology. However, the upstream signaling required for CIP2A-TOPBP1 foci formation was different. ATM activity is required for the formation of IR-induced CIP2A-TOPBP1 mitotic structures, but not for formation of APH-induced CIP2A-TOPBP1 mitotic structures. This requirement aligns with a role for MDC1 in TOPBP1 recruitment upon mitotic DSB generation^31,32,39^. Moreover, these data show that replication-mediated DNA lesions likely recruit CIP2A-TOPBP1 in a MDC1-independent fashion.

We also find a differential composition of CIP2A-TOPBP1 complexes upon IR compared to APH treatment. Our proteomic analysis uncovered that the SMX complex is a mitotic interactor of the CIP2A-TOPBP1 complex. In line with our proteomic results, APH-treated cells showed SMX complex localization to a large majority of mitotic CIP2A structures, whereas the majority of IR-induced mitotic CIP2A foci remained SMX-negative. This suggests that not all mitotic CIP2A-TOPBP1 structures are formed equally and that ancillary factors or post-translational modifications, instructed by the nature of the DNA lesion will determine the composition of these mitotic DNA response complexes.

SMX complex members were previously shown to resolve joint DNA molecules in a cell cycle-dependent fashion. Specifically, Cyclin-dependent kinase 1 (CDK1) and Polo-like kinase-1 (PLK1) promote the activity of MUS81-EME1^49–54^ and promote assembly of the SMX complex^55^. In addition, studies in yeast showed that the SLX4 scaffold of the SMX complex associates with the TOPBP1 orthologue Dpb11, promoted by CDK1-mediated phosphorylation of Thr1260^56^. Our data show that CIP2A is essential for the recruitment of SLX4 and associated SMX components during mitosis. The observed complete loss of mitotic TOPBP1 recruitment upon disruption of CIP2A likely prevents SLX4 and SMX recruitment in *CIP2A*^-/-^ cells. Similarly, a CIP2A mutant lacking the C-terminal 29 amino acids (CIP2A-ΔC) supports initial recruitment of TOPBP1, but does not form higher-order CIP2A-TOPBP1 structures and is defective for SLX4 recruitment. Although the exact function of the unstructured C-terminus of CIP2A remains elusive, based on our data one could hypothesize that the C-terminus facilitates filament formation between CIP2A homo-dimers. Mutation of the PLK1 phosphorylation site S904 only partially phenocopied the CIP2A-ΔC mutant, suggesting that other functional domains reside in the C-terminus. Previously reported structural studies using cryo-electron microscopy (cryo-EM) involved a truncated CIP2A construct (1-560) due to instability of the full-length protein^47^, which precluded analysis and positioning of the C-terminal domain. However, this truncation may have compromised the CIP2A-TOPBP1 interaction. Cryo-EM analysis of full-length CIP2A in combination with TOPBP1 is warranted to uncover insights into the formation and architecture of this complex.

Based on the different CIP2A-TOPBP1 structures we observed at the various stages of mitosis, it is tempting to speculate that the loop-containing CIP2A-TOPBP1 structures reflect repair intermediates, involving chromatin alterations to allow processing of the DNA lesions. Whether the filamentous CIP2A-TOPBP1 structures we predominantly observed in later stages of mitosis reflect ultrafine bridges (UFBs) is unclear. Interestingly, TOPBP1 was previously demonstrated to localize to UFBs in DT40 cells^57,58^.

The observation that the abundance of CIP2A-TOPBP1 structures gradually decreases during the course of mitotic progression already suggested that CIP2A-TOPBP1 is not only involved in tethering DNA ends, but also functions in processing of DNA lesions. One form of mitotic DNA lesion processing is MiDAS, which we show does not require CIP2A. This observation contributes to an ongoing debate regarding the molecular requirements for MiDAS. In RPE1 *TP53^-/-^* cells, our data align with a previously reported study that MUS81 is not required for MiDAS induced by APH^46^. Conversely, our previous work demonstrated that MUS81 is required for MiDAS induced by Cyclin E1 overexpression in RPE1 *TP53^-/-^* cells^18^. Similarly, short-term depletion of MUS81, SLX4 or EME1 in U2OS cells was reported to impair MiDAS^21^. These contradictory results about the role of the SMX complex in MiDAS ^18,21,46^ suggest that MiDAS has context-dependent requirements. Potentially, complex DNA lesions such as those involving oncogene-induced R-loops, may require additional processing when compared to under-replicated DNA induced by APH in this study.

Our finding that CIP2A is not required for MiDAS in RPE1 *TP53^-/-^*cells also aligns with reported synthetic lethal interactions^31^. A strong synthetic lethal interaction was observed between *CIP2A* and *BRCA1/2* in RPE1 *TP53^-/-^* cells. However, *RAD52,* which is essential for MiDAS in RPE1 *TP53^-/-^* cells, does not show profound synthetic lethality with *BRCA1/2* in these cells^31^. Combined, these findings argue against a major role for MiDAS underlying the synthetic lethality between *CIP2A* and *BRCA1/2*.

The mechanism by which SMX-mediated DNA breaks at under-replicated DNA are repaired during mitosis remains largely unclear. However, recent studies have demonstrated a role for CIP2A-TOPBP1 in the regulation of microhomology-mediated end joining (MMEJ) during mitosis via RHINO and POLQ^59,60^. Combined with our finding that the CIP2A-TOPBP1 complex recruits the SMX complex, a model emerges in which SMX-mediated breaks in under-replicated regions are repaired by POLQ-mediated microhomology-mediated end-joining^61^.

## Acknowledgements

The work described in this study was financially supported by the Dutch Cancer Society (grant #12911 to M.A.T.M.v.V) and the Netherlands Organization for Scientific Research (NWO VICI #09150182110019 to M.A.T.M.v.V, OCENW.M #21.106 to R.V.).

We thank dr. Manual Stucki, dr. Andrew Blackford, dr. Wojciech Niedzwiedz and dr. Peter Martin for constructive discussions and discussing unpublished work.

## Data availability

The mass spectrometry proteomics data have been deposited to the ProteomeXchange Consortium via the PRIDE [1] partner repository with the dataset identifier PXD059881.

## Code availability

The interactive Wiener Filter used for deconvolution of STED images using Jupyter Notebook has been deposited to Zenodo. This code can be found at https://doi.org/10.5281/zenodo.15075205.

## Methods

### Cell culture

All cell lines were grown in a humified incubator at 37 degrees Celsius in 5% CO_2_ and 20% O_2_. Human hTERT immortalized retinal pigmented epithelial RPE1 cells and human embryonic kidney HEK293T cells were obtained via the American Type Culture Collection (ATCC). DLD1 wildtype and DLD1 BRCA2^-/-^ human colorectal adenocarcinoma cells were from Horizon (Cambridge, UK). RPE1 *TP53*^-/-^ *PAC^-/-^* cells and RPE1 TP53^-/-^ *PAC*^-/-^ *BRCA1*^-/-^ cells were previously described^62^. HEK293T, MDA-MB231 and RPE1 cells were cultured in Dulbecco’s modified Eagle medium (DMEM, Thermofisher) complemented with 10% (v/v) fetal calf serum (FCS), 1% penicillin and 1% streptomycin (Gibco). HCC38, BT549 and DLD1 cells were cultured in Roswell Park Memorial Institute medium (RPMI, Thermofisher), complemented with 10% (v/v) FCS, 1% penicillin and 1% streptomycin (Gibco).

### Mutagenesis

RPE1 cells harboring a *TP53* mutation in exon 4 were described previously^18,63^. To generate the *CIP2A* knockout cell lines, a sgRNA (5’-ATCGGTTTGCTGTCTCAACT-3’) targeting exon 3 was cloned into pSpCas9(BB)-2A-GFP (px458) kindly provided by Feng Zhang (Addgene #48138)^64^. Upon transient transfection using Fugene HD transfection, GFP-positive cells were sorted with a Sony SH800S cell sorter at 48 h after transfection. Lack of *CIP2A* expression in two RPE1 *TP53^-/-^* clones (*CIP2A*^-/-^ cl#1 and cl#2), one MDA-MB321 clone and one HCC38 clone was confirmed by Western blot and immunofluorescence. *CIP2A* mutations in the two RPE1 *TP53^-/-^*clones were also confirmed with Sanger sequencing (cl#1: −1 deletion and +1 insertion, cl#2: +1 insertion).

### Cloning

Full length CIP2A was cloned from pcDNA3.1/CIP2A(1-905) WT V5 His (Addgene #119287), which was a gift from Jukka Westermarck^47^, into retroviral pMSCV-blast which was a gift from David Mu (Addgene #75085)^65^, and subsequently different CIP2A mutations were generated in this pMSCV plasmid.

To establish cell lines expressing doxycycline inducible short-hairpin RNAs (shRNAs) against scrambled (5’-CAACAAGATGAAGAGCACCAA-3’), luciferase (5’-AGAGCTGTTTCTGAGGAGCC-3’), *BRCA2* (5’-AACAACAATTACGAACCAAACTT-3’) and *CIP2A* (5’-GCTAGTAGACAGAGAACATAA-3’), DNA oligos were cloned as previously described into Tet-pLKO-puro vector (Addgene #21915)^66^. This Tet-pLKO-puro vector was a gift from Dmitri Wiederschain^67^.

### Short interfering RNA interference

Cells were seeded 24 h before transfection with short interfering RNAs (siRNAs) for the negative control (SR-CL-000-005, Eurogentec), siMUS81 (#1: CAGCCCUGGUGGAUCGAUA and #2: CAUUAAGUGUGGGCGUCUA), siCIP2A (GACAACUGUCAAGUGUACCACUCUU) and siSLX4 (AAACGUGAAUGAAGCAGAA). Cells were once or twice transfected with siRNAs using Oligofectamine^TM^ transfection reagent (Thermofisher) or Lipofectamine RNAiMAX (Invitrogen) as previously described^18,62^. Cells were either harvested for Western blot or fixed for immunofluorescence 48 h after siRNA transfection.

### Immunofluorescence microscopy

Cells were seeded on glass coverslips in 6-well or 12-well plates at 48 h prior to indicted treatments. Cells were fixed for 15 min using 2% paraformaldehyde in PBS. After fixation, cells were permeabilized with 0.5% Triton-X in PBS for 10 min. For MiDAS, EdU Click-IT reaction was performed for 30 min at room temperature according to protocol Click-IT^TM^ EdU Cell Proliferation Kit for Imaging (Invitrogen). Further details are provided in Supplementary Methods.

### STED microscopy

Detailed methodology is provided in the Supplementary Methods section. STED microscopy slide preparation was performed largely as previously described for fixed HEK293T cells^68^. STED imaging of fixed samples was performed on a STED microscope (Abberior Expert Line) with a 100× oil immersion objective (Olympus Objective UPlanSApo 100×/1.40 oil). During imaging, z-stacks of cells were made at confocal resolution and all CIP2A foci in each cell were subsequently selected manually and individually imaged at STED resolution using a custom Python script adapted from Mol and Vlijm^69^. Whereas other microscopy techniques often require reference measurements and corrections to determine the relative localization between different color channels, the use of a single STED depletion donut for both excitation colors counteracts potential wavelength dependent optical effects, resulting in almost perfect coalignment of both excitation channels^70^.

### Mass spectrometry

For on-bead digestion of immunoprecipitated proteins, bead mixtures were subjected to cysteine reduction followed by alkylation, and trypsin (Promega) digestion. Details are provided in the Supplementary Methods section. The mass spectrometry proteomics data have been deposited to the ProteomeXchange Consortium via the PRIDE partner repository with the dataset identifier PXD059881.

### In silico protein prediction

To predict the structure of CIP2A and TOPBP1, we used Alphafold Multimer (AF2 multimer v3)^71,72^ run on ColabFold^73^ with standard settings (pair mode: unpaired, paired, 5 models with 3 or 10 recycles). Models were ranked according to their predicted template modelling (pTM) scores and top-ranked models were analyzed and visualized in ChimeraX^74^. pAE plots were generated with ChimeraX^74^, and were annotated using Adobe Illustrator.

