## Supplemental Information for "CIP2A is required for mitotic recruitment of the SLX1/XPF/MUS81 tri-nuclease complex to replication stress-induced DNA lesions to maintain genome integrity"

This file includes:

Supplementary methods
Supplementary table legend
Supplementary Figure 1
Supplementary Figure 2
Supplementary Figure 3
Supplementary Figure 4
Supplementary Figure 5
Supplementary Figure 6
Supplementary Figure 7
Supplementary Figure 8

**Supplementary Methods**

##### *Viral transductions*

To establish CIP2A reconstituted RPE1 *TP53*$^{-/-}$, HEK293T cells were transfected with indicated pMSCV plasmids in combination of pRetro-VSV-G, pRetro-gag/pol and pAdvantage. Virus containing supernatant was harvested 48 h after transfection and filtered a 0.45 µm syringe filter. Virus-containing supernatant with 4 ug/mL of polybrene was subsequently used to infect target cells for 12 h. Transduced reconstituted cell lines were selected with 10 ug/mL blasticidin.

To generate doxycycline-inducible shRNA expressing cell lines, cells were transduced with indicated lentiviral pLKO shRNA plasmids. In brief, HEK293T cells were transfected with Tet-pLKO-puro vector with indicated shRNAs in combination with the lentiviral packaging plasmids pMD2.G and psPAX2. Virus containing supernatant was harvested 48 h and 56 h after transfection and filtered through a 0.45 µm syringe filter. Virus-containing supernatant with 4 ug/mL of polybrene was subsequently used to infect the target cells for 12 h. RPE1 *TP53*$^{-/-}$, RPE1 *TP53*$^{-/-}$ *CIP2A*$^{-/-}$ and *TP53*$^{-/-}$ *CIP2A*$^{-/-}$ cells expressing CIP2A mutants were transduced with indicated pLKO shRNAs and were selected with 15 ug/mL puromycin. BT549 cells transduced with pLKO shRNAs were selected with 1ug/mL puromycin. Cells were harvested for Western blot, seeded for clonogenic survival assays, or fixed for immunofluorescence 48 h after shRNA induction by addition of 1 ug/mL doxycycline.

##### *Immunofluorescence microscopy*

Cells were seeded on glass coverslips in 6-well or 12-well plates at 48 h prior to treatments. If indicated, cells were treated for 16 h with low dose APH (200 nM, Merck) before addition of RO-3306 (5µM, Axon Medchem) for 4 h to arrest cells at G2/M-border. RO-3306 was washed out from synchronized cells to allow entry into mitosis by washing once with pre-warmed PBS, followed by releasing the cells in complete medium. For analysis of IR-induced DNA damage foci in mitosis, cells were irradiated with 0.25 Gy 5 min after release using a Cesium$^{137}$ source and fixed 30 min after release. For micronuclei quantification, asynchronous cells were treated for 24 h with 200 nM APH before fixation or cells were treated with 3 Gy IR and recovered for 72 h before fixation. For MiDAS analysis, 20 uM EdU was added to complete medium during release. For ATM and ATR inhibition cells were pretreated 30 min before RO-3306 release with either 10 µM ATRi KU55933 (Axon Medchem) or 10 µM ATMi VE-821(MedChem Express), and subsequently released in presence of the inhibitors. Cells were fixed 25 min after release for 15 min using 2% paraformaldehyde in PBS. After fixation, cells were permeabilized with 0.5% Triton-X in PBS for 10 min. For MiDAS, EdU Click-IT reaction was performed for 30 min at room temperature according to protocol Click-IT™ EdU Cell Proliferation Kit for Imaging (Invitrogen). Cells were subsequently incubated for 1 h with blocking buffer containing 2% bovine serum albumin (BSA) and 0.05% Tween in PBS (PBT). Incubation with primary antibodies was performed overnight at 4° Celsius with indicated antibodies in PBT. The following antibodies were used: mouse anti-γH2AX (Millipore, 05-636, 1:400), rabbit anti-γH2AX (Cell Signaling, 9718, 1:400), mouse anti-CIP2A (Santa Cruz, sc-80659, 1:500), rabbit anti-CIP2A (Invitrogen, PA5-83469, 1:300), rabbit anti-TOPBP1 (Bethyl, A300-111A-M, 1:400), mouse anti-ERCC1 (Santa Cruz, sc-17809, 1:100), rabbit anti-XPF (Abcam, ab76948, 1:100), mouse anti-MUS81 (abcam, ab14387, 1:100), mouse anti-BTBD12 (SLX4) (Abnova, H000844644-B01P, 1:500), mouse anti-V5 (Invitrogen, R960-25, 1:400), rabbit anti-V5 (Cell signaling, 13202S, 1:1000). After washing the coverslips to remove residual primary antibodies, incubation with DAPI and the following secondary antibodies: 488 goat anti-mouse (Thermofisher, A11029, 1:500), 488 goat anti-rabbit (Thermofisher, A11008, 1:500), 488 donkey anti-rabbit (Thermofisher, A21206, 1:500), 647 goat anti-mouse (Thermofisher, A32728, 1:500), 647 donkey anti-rabbit (Thermofisher, A31573, 1:500), 647 donkey anti-mouse (1:500) in PBT was performed at room temperature. For micronuclei analysis, cells were only incubated with DAPI after permeabilization. Coverslips were extensively washed and subsequently mounted with Prolong Gold antifade reagents (Life Technologies). Images were acquired using Zeiss Axio Imager 2 using a 40x or 63x objective with the Zen 3.0 software (Zeiss) or on a Leica DM6000B microscope using a 10x or 40x objective with LAS-AF software (Leica), and analysis was performed using FIJI (ImageJ). V5 and TOPBP1 foci intensity were quantified using the ImageJ macro 'Foci-analyzer' (freely available at https://github.com/BioImaging-NKI/Foci-analyzer; created by Bram van den Broek, the Netherlands Cancer Institute, the Netherlands). Fluorescence signal intensities are depicted as arbitrary units, and are normalized by dividing the mean foci intensity per cell by the median intensity of RPE1 *TP53*$^{-/-}$ *CIP2A*$^{-/-}$ cells with reconstituted CIP2A-WT-V5 per individual experiment.

***STED microscopy***
For STED microscopy, cells were seeded, treated and fixed similar as described above for immunofluorescence microscopy, with the following exception: cells were fixed 30 min after RO-3306 washout to obtain cells at early mitotic stages, whereas cells were fixed at 60 min and 90 min after RO-3306 washout to obtain cells at later stages of mitosis. STED microscopy slide preparation was performed largely as previously described for fixed HEK293T cells[68]. Specifically, cells were incubated with 2% BSA, 10% goat serum, and 0.05% Tween as blocking buffer. This specific blocking buffer was also used for primary and secondary antibody incubations. Primary antibodies were used as earlier described, and the following secondary antibodies were used: 580 goat anti-mouse (Abberior STAR, ST580-1001, 1:80), 580 goat anti-rabbit (Abberior STAR, ST580-1002, 1:80), 635 goat-anti mouse (Abberior STAR, ST635-1001, 1:80), 635 goat anti-rabbit (Abberior STAR, ST635-1002, 1:80).

STED imaging of fixed samples was performed on a STED microscope (Abberior Expert Line) with a 100× oil immersion objective (Olympus Objective UPlanSApo 100×/1.40 oil). Laser alignments were optimized before imaging using a 0.1-µm bead sample (Invitrogen, TetraSpeck T7279). During imaging, z-stacks of cells were made at confocal resolution and all CIP2A foci in each cell were subsequently selected manually and individually imaged at STED resolution using a custom Python script adapted from Mol and Vlijm[69]. All STED images were acquired with a pixel size of 22 nm, a pixel dwell time of 20 µs and a 0.6 Airy Units (AU) pinhole, with image size varying based on structure size. For excitation, we used 40 MHz pulsed lasers with a wavelength of 561 nm (203 µW at laser head) and 640 nm (1 mW at laser head) and a continuous wave laser with a wavelength of 405 nm for DAPI excitation (confocal only). A 40 MHz pulsed laser (3.0 W) with a wavelength of 775 nm was used for depletion. Other microscopy techniques often require reference measurements and corrections to determine the relative localization between different color channels. On the contrary, the use of a single STED depletion donut for both excitation colors counteracts potential wavelength dependent optical effects, resulting in almost perfect coalignment of both excitation channels[70]. Detection filters were set to 699 ± 52 nm for the 640 channel (640 nm excitation) and 600 ± 31 nm for the 561 channel (561 nm excitation). The 640 and 561 channel were collected separately per line, except for EdU-CIP2A images, in which both channels were collected in separate images (first

640 nm then 561 nm channel) to prevent bleaching of the Alexa Fluor 647 dye during imaging of the 561 nm channel. The applied laser powers, dwell times, line steps and gating for STED imaging were optimized for the different samples and dyes: STAR 635 (0.9-1.5% excitation (640 nm), 28-35% STED, 12-24 lines, 0.75-1.2 ns gating delay, 9.0 ns gating width), STAR 580 (4.1-25.0% excitation (561 nm), 45-60% STED, 12 lines, 0.45-1.0 ns gating delay, 9.0 ns gating width). For EdU-CIP2A samples, the following settings were used: EdU/Alexa Fluor™ 647 (1.9-2.0% excitation (640 nm), 4% STED, 6 lines, gating off), CIP2A/STAR 580 (17.2-50.0% excitation (561 nm), 50% STED, 12 lines, gating off).

##### *STED image processing*

The STED images were, unless specified differently, deconvoluted using a custom-written Wiener filter (Mol, F. N., & Vlijm, R. (2025). Interactive Wiener Filter (v1.0). Zenodo. https://doi.org/10.5281/zenodo.15075206), with background subtraction of 2-50 counts (640 channel) or 2-30 counts (561 channel), STED/confocal ratio of 0.7-0.9 (640 channel) or 0.6-0.8 (561 channel), noise-to-signal ratio of 0.05-0.3 (640 channel) or 0.08-0.25 (561 channel) and point-spread functions of 26-35 nm (STED) and 350 nm (confocal) for the 640 channel or 30-35 nm (STED) and 300 nm (confocal) for the 561 channel. Due to the large spread in intensities, linear intensity scaling was applied separately for each image. The structure length was determined by multiplying the manually measured length in pixels by the pixel size of 22 nm. These structure size measurements were performed using FIJI (ImageJ). Line profiles were drawn in FIJI with a line width of 3 pixels and a segmented line with spline fit was used for curved line profiles.

##### *Co-immunoprecipitation*

RPE1 *TP53*$^{-/-}$ and RPE1 *TP53*$^{-/-}$ *CIP2A*$^{-/-}$ cl#1 cells were seeded in T175 flasks for co-immunoprecipitation (IP) of endogenous CIP2A in mitotic cells. For IP in irradiated conditions, cells were treated with 62.5 ng/mL nocodazole for 16 h prior to mitotic shake off to harvest mitotic cells. Mitotic cells were either left untreated (control) or treated with 5 Gy irradiation and recovered for 1 h before adding ice-cold MPER lysis buffer (Thermofisher), complemented with protease and phosphatase inhibitor cocktail (Thermofisher) and 5 units/mL benzonase (Santa Cruz) (‘complete lysis buffer’). For

IP in APH-treated cells, cells were either left untreated (control) or treated with 200 nM APH for 16 h prior to adding 5 μM RO-3306 for 4 h. APH treated cells were subsequently pre-treated 30 min before release with 5 mM hydroxyurea (HU). Synchronized cells were released from RO-3306 by washing once with pre-heated PBS, followed by RO-3306 wash-out using complete medium including 62.5 ng/mL nocodazole (control) or 62.5 ng/mL nocodazole and 5 mM HU (APH treated cells) to accumulate cells in mitosis. Mitotic shake off was performed at 1 h after release and cells were lysed with ice-cold complete lysis buffer.

Both APH and IR samples were tumbled end-over-end for 30 min at 4° Celsius after addition of complete lysis buffer, and subsequently centrifuged for 15 min at 12,000 rpm to clear whole cell lysates. The protein concentration of the supernatant was quantified using Pierce BCA Protein Assay Kit (Thermofisher). 1 ug of CIP2A antibody was added to the pre-cleared whole cell lysates for 1 h and tumbled end-over-end at 4 degrees Celsius. Subsequently, magnetic Dynabeads Protein G beads (Thermofisher) for immunoprecipitation were added to the lysates and incubated for another hour at 4° Celsius. Beads were precipitated using a magnet and were washed 3 times with low salt buffer (20 mM HEPES pH 7.5, 150 M NaCl in $dH_2O$), and a final wash with high salt buffer (20 mM HEPES pH 7.5, 496 M NaCl in $dH_2O$). Protein samples for western blot analysis were eluted from the beads after this final high salt wash step by boiling the beads in 2x SDS-sample buffer with beta-mercaptoethanol, whereas samples for mass spectrometry were dissolved in denaturing buffer (8M Urea 100 mM Tris pH8,0) and snap frozen.

***Mass spectrometry***

For on-bead digestion of immunoprecipitated proteins, bead mixtures were subjected to cysteine reduction (1.5 mM dithiothreitol for 15 min at 30°C), followed by alkylation (7.5 mM iodoacetamide, 30 min at RT). Bead mixture was diluted to 2M Urea with 100 mM ammonium bicarbonate and digested with 100 ng trypsin (Promega) overnight at 37°C. After bead removal, peptides were acidified with 0.1% formic acid, extracted by solid phase extraction with C18 cartridges (Gracepure SPE C18-Aq), and dried by SpeedVac (Thermofischer). Lastly, extracted peptides were resuspended in 0.1% formic acid and analyzed in triplicate with a Exploris480 Orbitrap mass spectrometer (Thermofischer) setup for data independent acquisition (DIA). Acquired spectra were analyzed using Spectronaut v17.6 (Biognosys), and data was processed using

Perseus v1.6.15 (MaxQuant). Three technical replicate measurements were performed per sample. Statistical analysis was conducted in Perseus using two-tailed unpaired t-test. The mass spectrometry proteomics data have been deposited to the ProteomeXchange Consortium via the PRIDE partner repository with the dataset identifier PXD059881.

***Western blot***

Cells were lysed with MPER lysis buffer (Thermofisher), complemented with protease and phosphatase inhibitor cocktail (Thermofisher) in presence or absence of benzonase (100 units/mL, Merck) and EDTA (5 mM, Thermofisher). Protein concentration was measured using the Pierce™ BCA Protein Assay Kit (Thermofisher). Proteins were subsequently separated using SDS-polyacrylamide gels, and transferred to PVDF membrane (conventional transfer: Immobilon, turbo transfer: Bio-Rad). Membranes were blocked with 5% skimmed milk (Sigma) in Tris-buffered saline (TBS) with 0,05% Tween (TBS-T) or 3% BSA (Sigma) in TBS-T. Membranes were incubated with the following antibodies: mouse anti-CIP2A (Santa Cruz, sc-80659, 1:500), rabbit anti-TOPBP1 (Bethyl, A300-111A-M, 1:1000), rabbit anti-γH2AX (Cell signaling, 9718, 1:1000), mouse anti-Actin (MpBiomedicals, 69100, 1:10.000), HRP conjugated Beta-Actin (Proteintech, HRP-60008, 1:10.000), rabbit anti-GAPDH (Abcam, ab128915, 1:1000), rabbit anti-MDC1 (Abcam, ab11171, 1:1000), mouse anti-MUS81 (Abcam, ab14387, 1:1000), rabbit anti-Vinculin (Abcam, ab129002, 1:5.000), mouse anti-BTBD12 (SLX4) (Abnova, H00084464-B01P, 1:1000), mouse anti-BRCA2 (Calbiochem, OP95, 1:1000), mouse anti-HSP90α/β (F-8) (Santa Cruz, sc-13119. 1:10.000), rabbit α-tubulin (Cell signaling, 2125, 1:1000) overnight at 4° Celsius. Membranes were extensively washed and incubated with the following horse-radish peroxidase-conjugated secondary antibodies: HRP-conjugated rabbit anti-mouse (DAKO, 1:5000) and HRP-conjugated goat anti-rabbit (DAKO, 1:5000) for 1 h at room temperature and visualized using Lumi-light or SuperSignal West Femto Maximum Sensitivity Substrate (Thermofisher). Images were acquired with a ChemiDoc MP imaging system (Bio-Rad).

**Clonogenic survival assays**

RPE1 cells with either shLUC or shBRCA2 were pretreated for 48h with 1ug/mL doxycycline before seeding 100 or 200 into a 6-well plate. Fresh doxycycline was added during seeding, and medium was not refreshed during the entire experiment. Cells were fixed with methanol and stained with staining buffer (50% methanol, 29,95% water, 20% acetic acid and 0.05% Coomassie Brilliant Blue) 7-9 days after seeding. Colonies were imaged using an EliSpot reader (Alpha Diagnostics International) with vSpot Spectrum software. The number of colonies were manually counted.

**Supplementary Tables:**

**Supplementary Table 1: Mass spectrometry data.**

Data underlying Figures 2A, B and Supplemental Figure 2B, C are provided here.

**Supplemental Figures**

**Supplement figure 1: Different types of mitotic DNA damage recruit CIP2A-TOPBP1**

**(A)** Quantification of CIP2A foci colocalizing with gH2AX foci in RPE1 *TP53*$^{-/-}$ cells, either untreated or after treatment with APH (200 nM, 20 h) or IR (0,25 Gy). Data from one experiment with n>30 cells per experimental condition are shown. **(B)** Western blot analysis of CIP2A in parental RPE1 *TP53*$^{-/-}$ cells or *CIP2A*$^{-/-}$ clones. Lysates were immunoblotted for indicated proteins. **(C)** Representative images of parental RPE1 *TP53*$^{-/-}$ or *CIP2A*$^{-/-}$ cl#1 cells, stained for DAPI (blue), CIP2A (green) and TOPBP1 (red), in either untreated conditions, or after treatment with APH (200 nM, 20 h) or IR (0,25 Gy). Scale bar represents 10 µm. **(D)** Quantification of TOPBP1 foci per mitotic cell for cells treated as described in panel C. The individual values and medians of one experiment with n≥25 cells per experimental condition are shown. **(E)** Representative images of RPE1 *TP53*$^{-/-}$ cells and two *CIP2A*$^{-/-}$ clones treated with APH (200 nM, 24 h) and stained with DAPI. Scale bar represents 10 µm. Arrows indicate micronuclei. **(F)** Quantification of CIP2A foci per mitotic cell in RPE1 *TP53*$^{-/-}$ cells, either left untreated or treated with APH (200 nM, 20 h), IR (0,25 Gy), ATRi (VE-821, 10 µM) and/or ATMi (KU55933, 10 µM). Individual values and medians of two experiments with n>25 cells per experimental condition are shown. **(G, H)** Raw images of the Wiener-deconvolved STED data shown in Fig. 1D, E. **(I)** Quantification of CIP2A structures colocalizing with TOPBP1 as observed by STED microscopy. The bar represents the mean and SD of three experiments, n represents the total number of structures analyzed per treatment in three experiments. **(J)** Representative images of measurements of the longest axis of unstructured, loop-containing and filamentous CIP2A-TOPBP1 complexes. Arrows indicate the length measurements used in Fig. 1I for the CIP2A structures shown in Fig. 1D.

**Supplement figure 2: Replication stress induces CIP2A-TOPBP1-mediated SLX4 recruitment, facilitating the assembly of the SLX4-ERC1-XPF-MUS81 complex.**

**(A)** Western blot analysis of endogenous CIP2A co-immunoprecipitations in mitotic parental RPE1 *TP53*$^{-/-}$ cells and *CIP2A*$^{-/-}$ cl#1 cells after IR (5 Gy). Lysates were immunoblotted for indicates proteins. **(B)** Mass spectrometry analysis of mitotic endogenous CIP2A co-immunoprecipitations in untreated parental RPE1 *TP53*$^{-/-}$ versus untreated *CIP2A*$^{-/-}$ cl#1 cells as controls for co-immunoprecipitations from IR-treated cells. Proteins indicated in red are enriched after IR and APH treatment. **(C)** Mass spectrometry analysis of mitotic endogenous CIP2A co-immunoprecipitation in untreated RPE1 *TP53*$^{-/-}$ versus untreated *CIP2A*$^{-/-}$ cl#1 cells as controls for co-immunoprecipitation from APH-treated cells. Proteins indicated in red are enriched after IR or APH treatment. **(D)** Quantification of CIP2A foci and colocalizing CIP2A/XPF foci per mitotic cell in untreated RPE1 *TP53*$^{-/-}$ or in RPE1 *TP53*$^{-/-}$ cells treated with APH (200 nM, 20 h) or IR (0,25 Gy). Individual values and medians are plotted of one experiment with n>35 cells per experimental condition. **(E)** Western blot analysis of RPE1 *TP53*$^{-/-}$ cells transfected with siCTRL or siSLX4. Lysates were immunoblotted for indicated proteins. **(F)** Representative images of RPE1 *TP53*$^{-/-}$ cells stained for DAPI (blue), CIP2A (green) and SLX4 (red) treated with APH (200 nM, 20 h) transfected with either siCTRL or siSLX4. Scale bar represents 10 µm. **(G)** Representative images of RPE1 *TP53*$^{-/-}$ cells stained for DAPI (blue), CIP2A (green) and MUS81 (red) treated with APH (200 nM, 20 h) transfected with either siCTRL or siSLX4. Scale bar represents 10 µm. **(H)** Quantification of CIP2A foci per mitotic cell for cells treated as described in panel F. Individual values, medians and interquartile range of two experiments with n≥29 cells per experimental condition are plotted. **(I)** Quantification of SLX4 foci per mitotic cell for cells treated as described in panel F. Individual values, medians and interquartile range of two experiments with n≥29 cells per experimental condition are plotted. **(J)** Quantification of MUS81 foci per mitotic cell for cells treated as described in panel G. Individual values, median and interquartile range of two experiments with n>30 cells per experimental condition are plotted.

**Supplement figure 3: Raw STED data**

**(A-E)** Raw images of the Wiener-deconvolved STED data shown in Fig. 3A-C, G, I

**Supplement figure 4: Line profiles analysis of STED images.**

**(A-C)** Line profiles were drawn on dual-color STED images of CIP2A (green) together with SLX4, ERCC1 or MUS81 (red) in panel A, or yH2AX and EdU (red) in panels B, C for RPE1 *TP53*$^{-/-}$ cells, treated as in Figure 3. For each dual-color STED image, 2-3 lines are shown and the corresponding intensity profiles are shown on the right. STED images were processed only with mild background subtraction. Scale bar represents 500 nm.

**Supplement figure 5: Western blot verification of CIP2A loss or MUS81 depletion in multiple cell lines.**

**(A)** Western blot analysis of parental RPE1 *TP53*$^{-/-}$ cells, *CIP2A*$^{-/-}$ cl#1 cells, and *CIP2A*$^{-/-}$ cl#1 cells reconstituted with indicated CIP2A mutants. Lysates were immunoblotted for indicated proteins. **(B)** Western blot analysis of RPE1 *TP53*$^{-/-}$ cells transfected with indicated siRNAs. Lysates were immunoblotted for indicated proteins. Asterisk indicates an aspecific band. Quantification of EdU foci per mitotic cell in RPE1 *TP53*$^{-/-}$ cells transfected with indicated siRNAs. Individual values, medians and interquartile range of two experiments with n≥29 cells per experimental condition are plotted. **(C)** Western blot analysis of CIP2A in parental MDA-MB-231 cells and MDA-MB-231 *CIP2A*$^{-/-}$ cells. Lysates were immunoblotted for indicated proteins. Quantification of CIP2A foci per mitotic cell in parental MDA-MB-231 cells and MDA-MB-231 *CIP2A*$^{-/-}$ cells treated with APH (200 nM, 20 h). **(D)** Western blot analysis of CIP2A in parental HCC38 cells and HCC38 *CIP2A*$^{-/-}$ cells. Lysates were immunoblotted for indicated proteins. **(E)** Western blot analysis of CIP2A in untreated or doxycycline-treated BT549 with doxycycline-inducible scramble (shSCR) or CIP2A (shCIP2A) shRNA. Lysates were immunoblotted for indicated proteins.

**Supplement figure 6: Data supporting main figure 5A-C**

Plots with predicted Aligment Erros (pAE) for predictions of **(A)** TOPBP1 1-1522 with 2 copies of CIP2A 1-905 and **(B)** 2 copies of CIP2A 1-876 with 2 copies of TOPBP1 755-860. Low pAE scores indicated proximities with high confidence. Structural features apparent from this analysis include i) an extensive and high confidence CIP2A:CIP2A coiled coil interaction interface mediated by CIP2A's C-terminal region, ii) the putative interaction of TOPBP1 755-798 to the CIP2A coiled coil, and iii) the putative interaction of TOPBP1 830-849 with the globular N-terminal region of CIP2A. **(C)** Mapping of posttranslational modification (phosphosite plus) on the sequences of TOPBP1 and CIP2A. Modifications detected in 8 or more studies are indicated.

**Supplement figure 7: C-terminal domain of CIP2A is required for functional CIP2A-TOPBP1 complex and MUS81 recruitment**

**(A)** Representative images of *CIP2A*$^{-/-}$ cl#1 reconstituted with indicated CIP2A mutants stained for DAPI (blue), V5 (green) and TOPBP1 (red) treated with APH (200 nM, 20 h). Scale bar represents 10 µm. **(B)** Quantification of mean V5 foci and mean TOPBP1 foci intensity per mitotic cell for cells treated as described in panel B normalized to median foci intensity of *CIP2A*$^{-/-}$ cells reconstituted with CIP2A-WT. Individual values, median and interquartile range of three experiments with n>30 cells per experimental condition are plotted. P-values were calculated using ordinary one-way ANOVA with Dunnett's multiple comparison test on the medians per experiment. **(C)** Representative images of *CIP2A*$^{-/-}$ cl#1 reconstituted with either CIP2A-WT, CIP2A-S904A or CIP2A-ΔC stained for DAPI (blue), V5 (red) and MUS81 (green) treated with APH (200 nM, 20 h). Scale bar represents 10 µm. **(D)** Raw images of the Wiener-deconvolved STED data shown in Fig. 5I. **(E)** Raw images of the Wiener-deconvolved STED data shown in Fig. 5J.

**Supplement figure 8: Western blot verification of BRCA2, CIP2A and SLX4 depletion**

**(A)** Western blot analysis of either untreated or doxycycline-treated RPE1 *TP53*$^{-/-}$ cells, *CIP2A*$^{-/-}$ cl#1 cells and *CIP2A*$^{-/-}$ cl#1 cells reconstituted with indicated CIP2A mutants with doxycycline-inducible luciferase (shLUC) andBRCA2 (shBRCA2). Lysates were immunoblotted for indicated proteins. **(B)** Western blot analysis of DLD1 WT and DLD1 *BRCA2*$^{-/-}$ cells transfected with indicated siRNAs. Lysates were immunoblotted for indicated proteins.

### Supplemental Figure 1

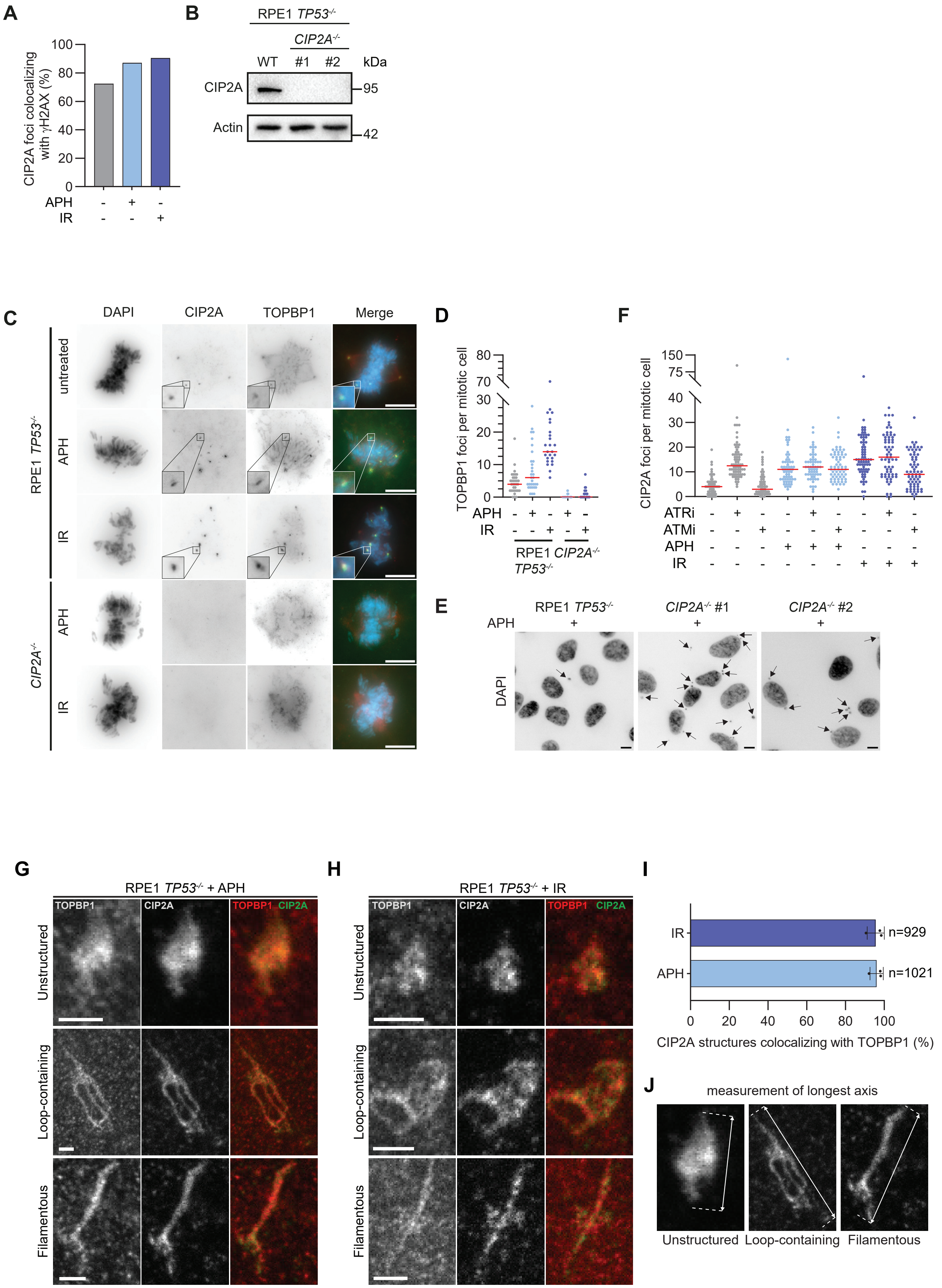

### Supplemental Figure 2

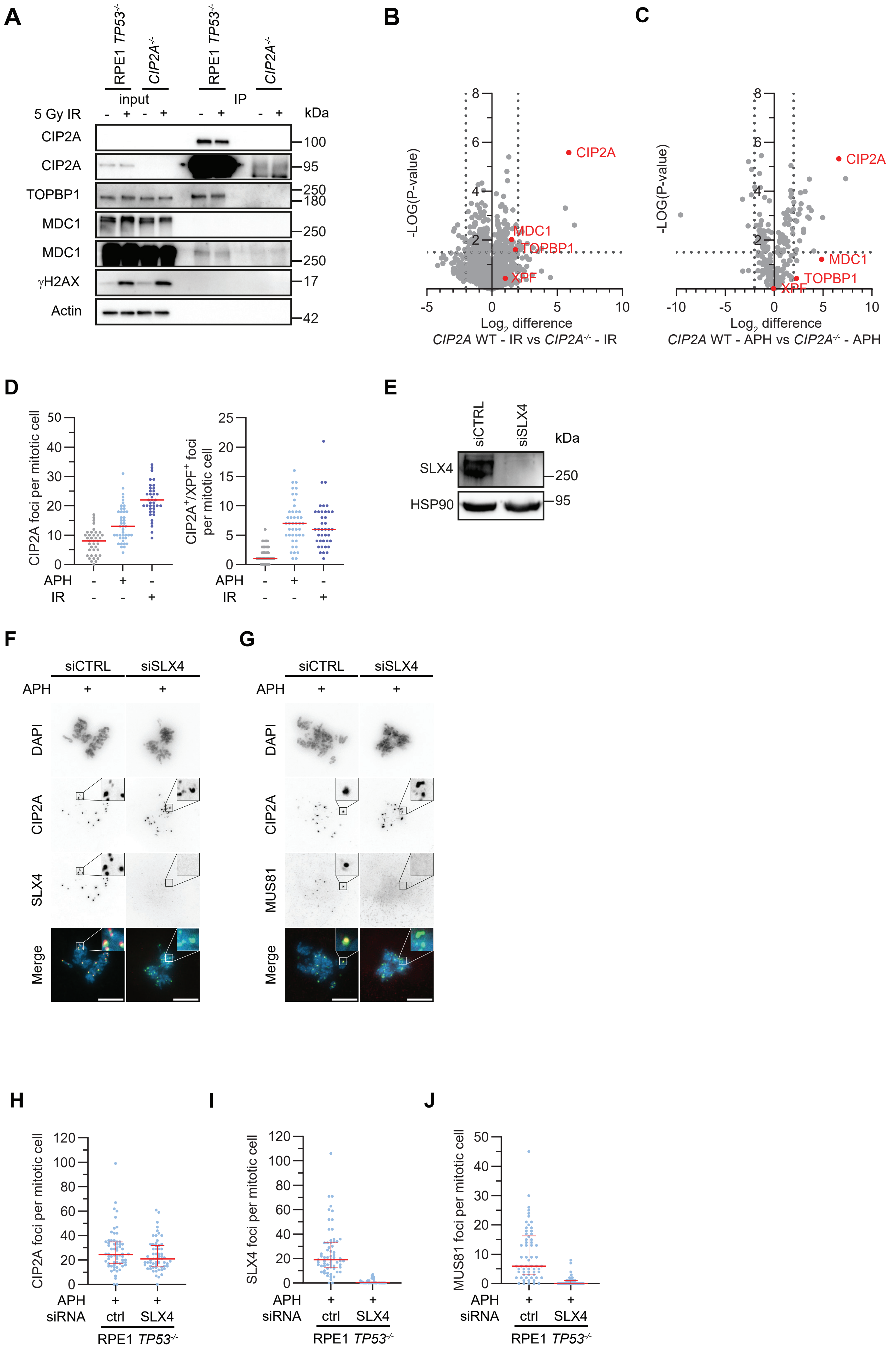

### Supplemental Figure 3

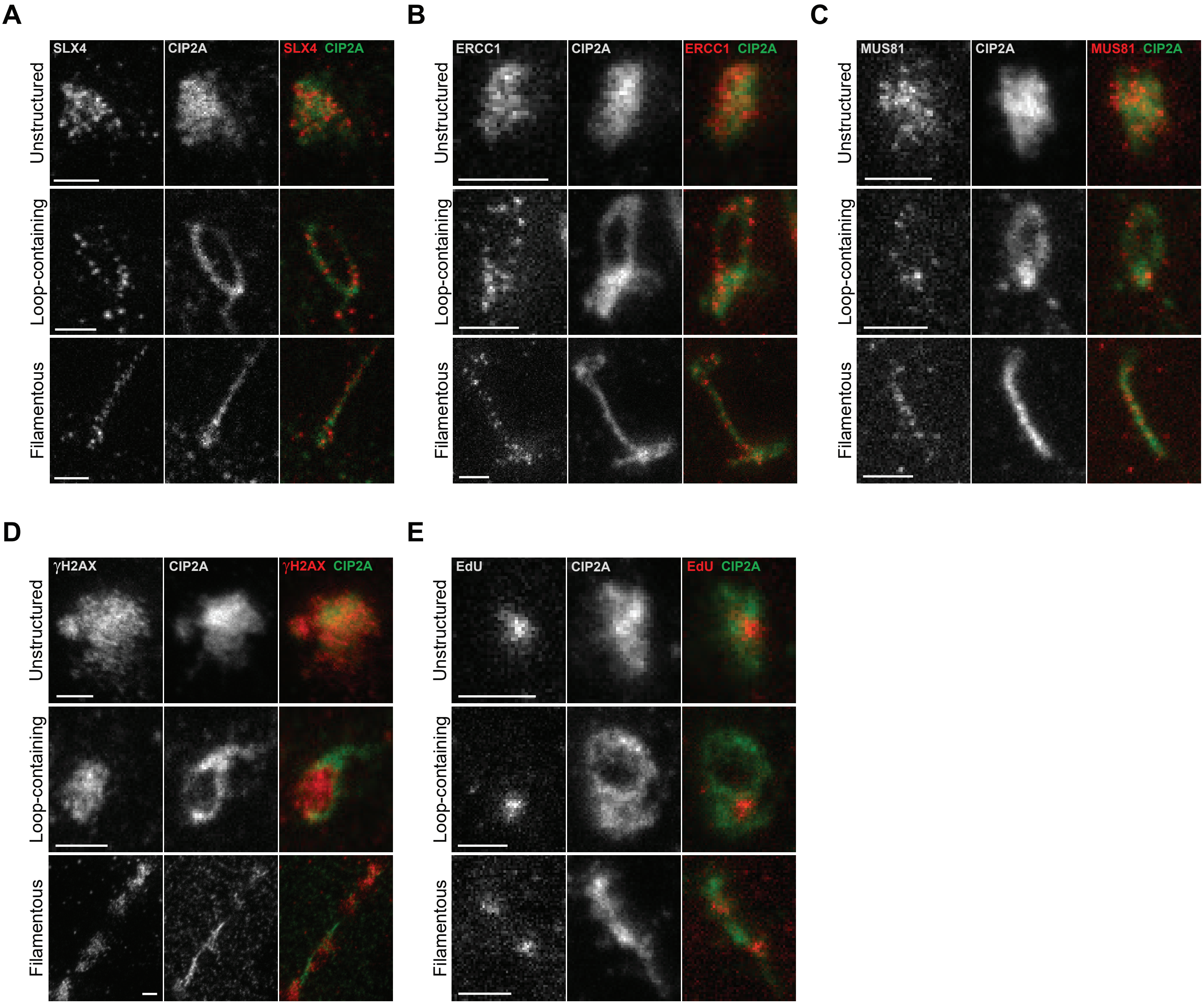

#### Supplemental Figure 4

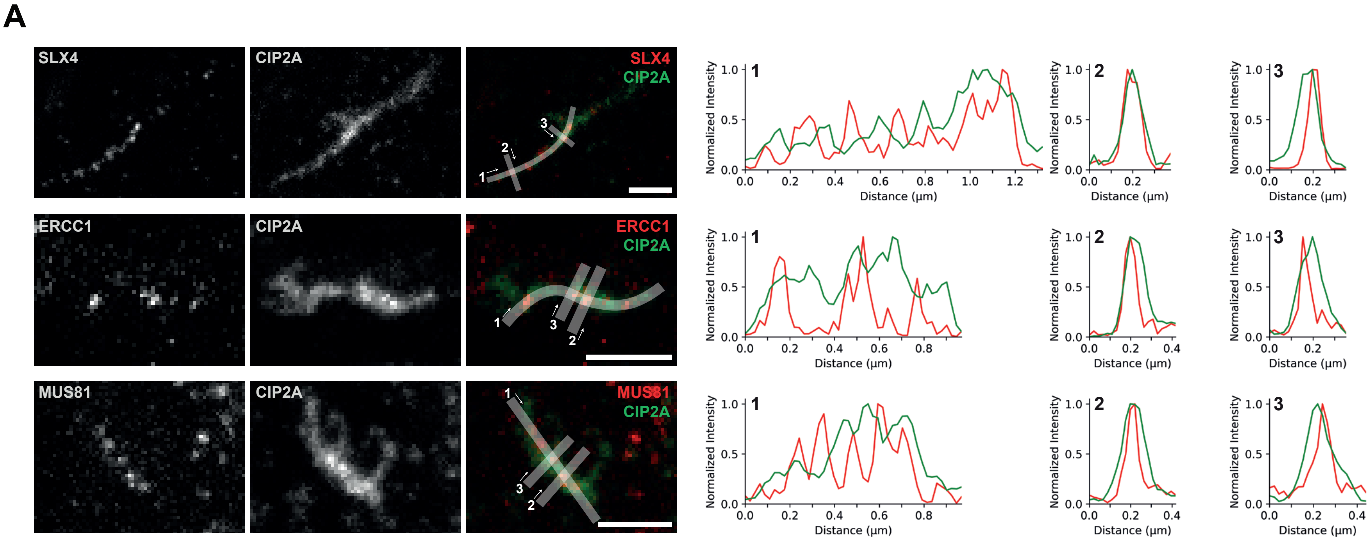


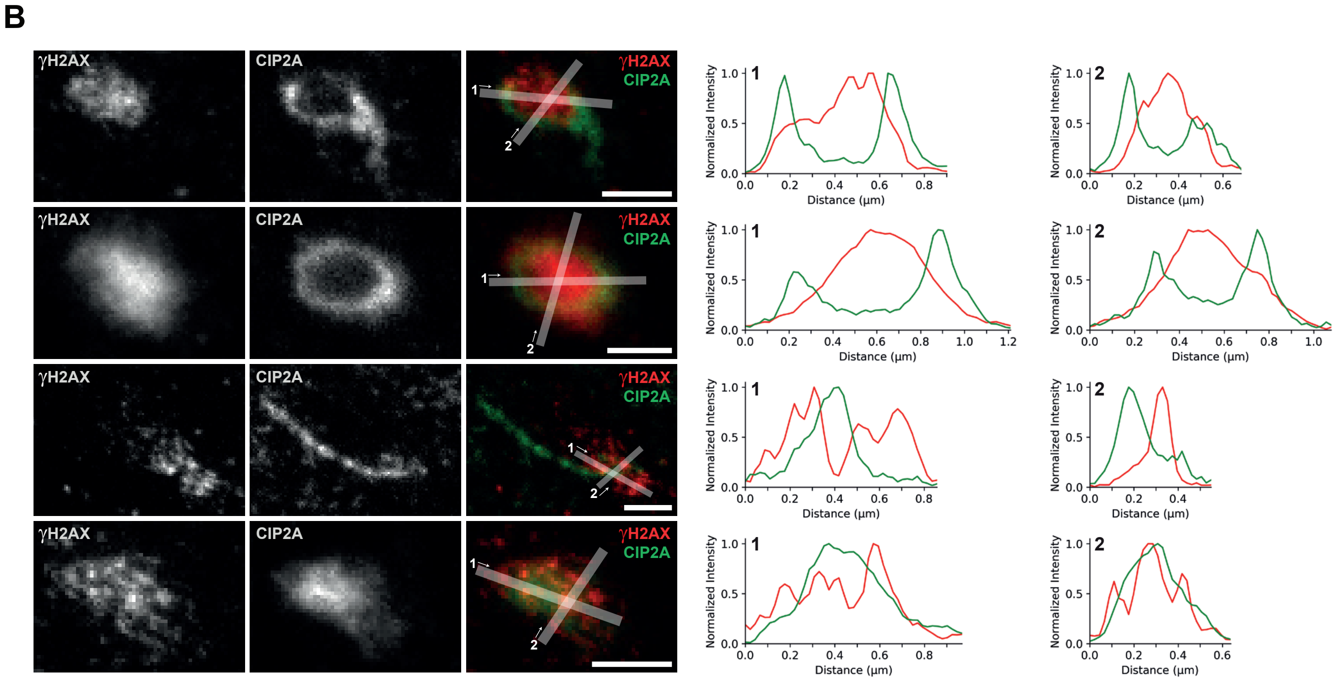


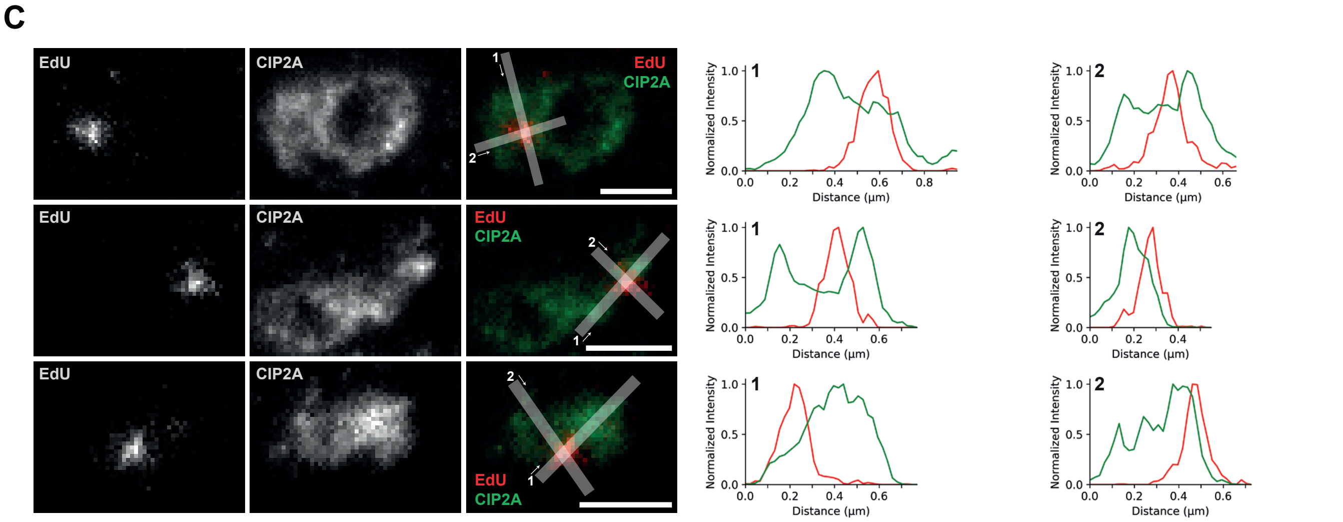

### Supplemental Figure 5

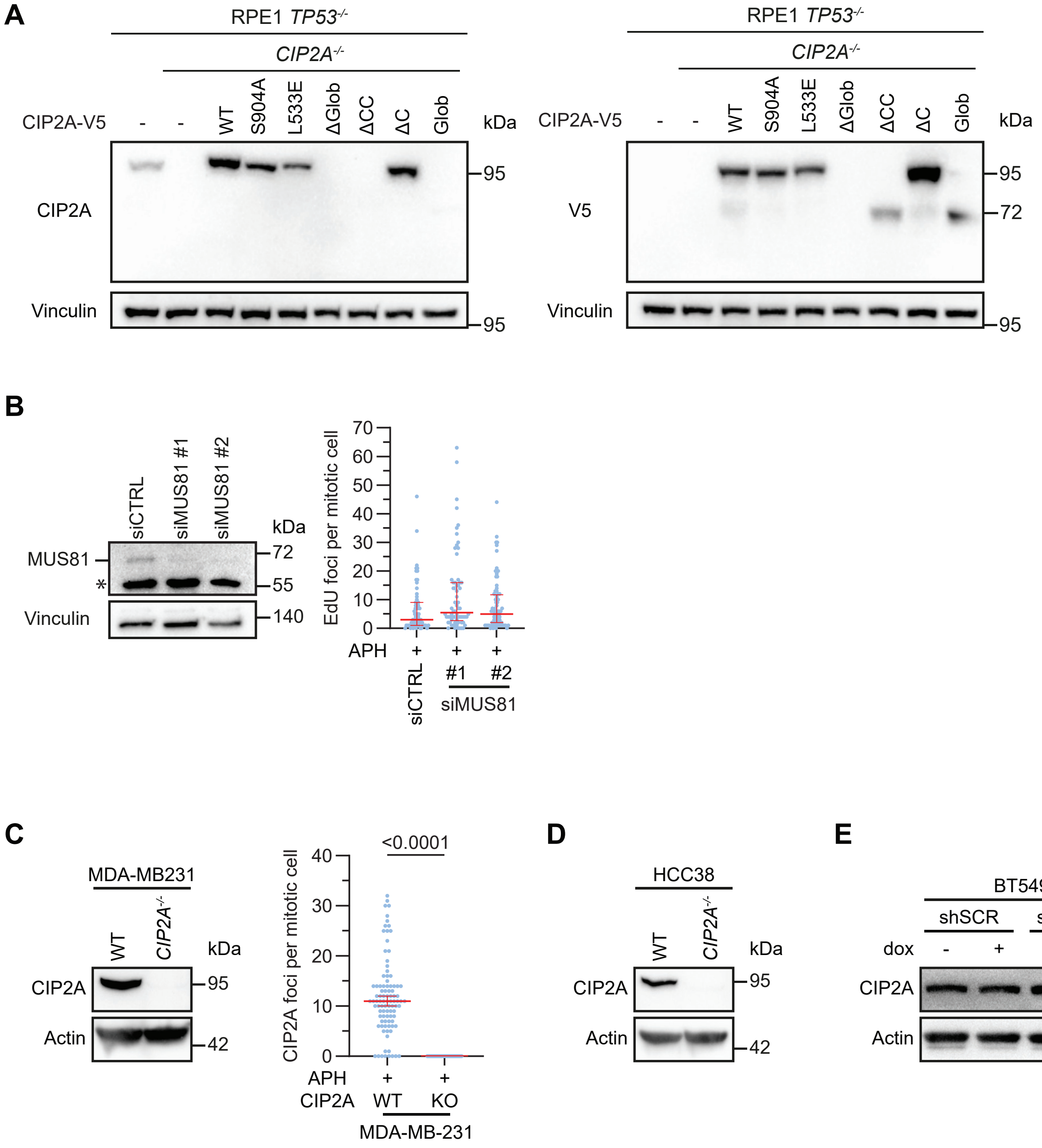

### Supplemental Figure 6

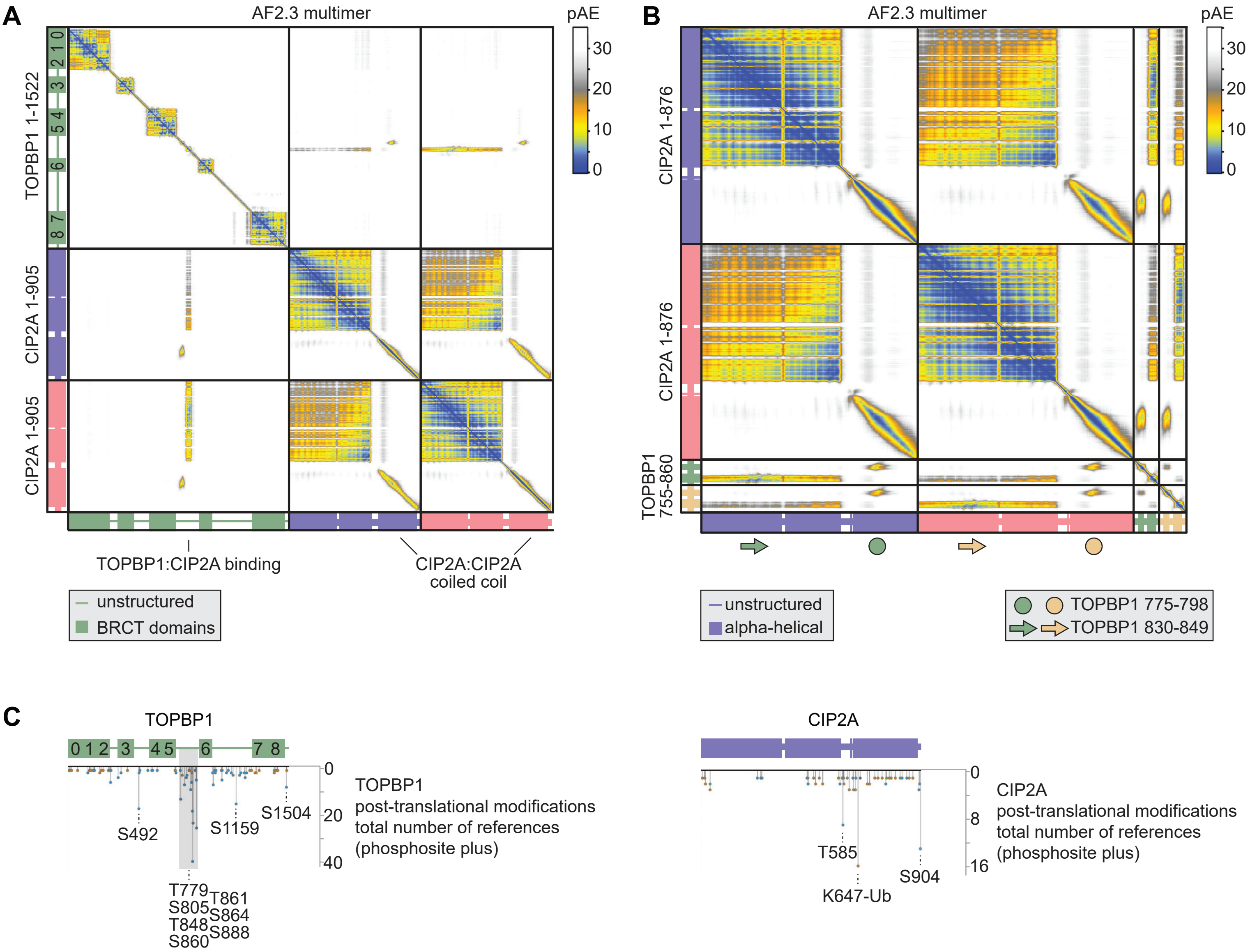

### Supplemental Figure 7

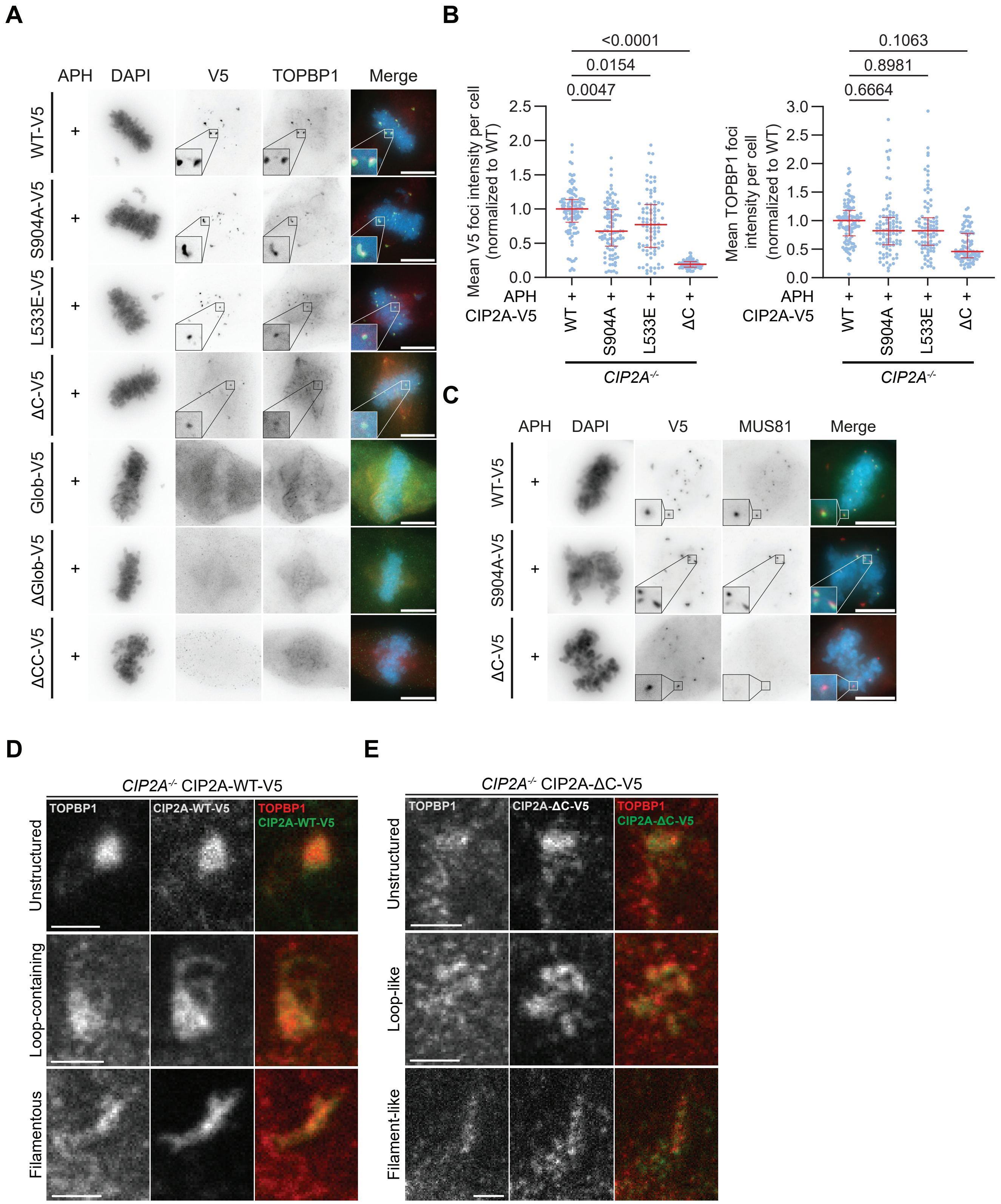

**Supplemental Figure 8**

**A**

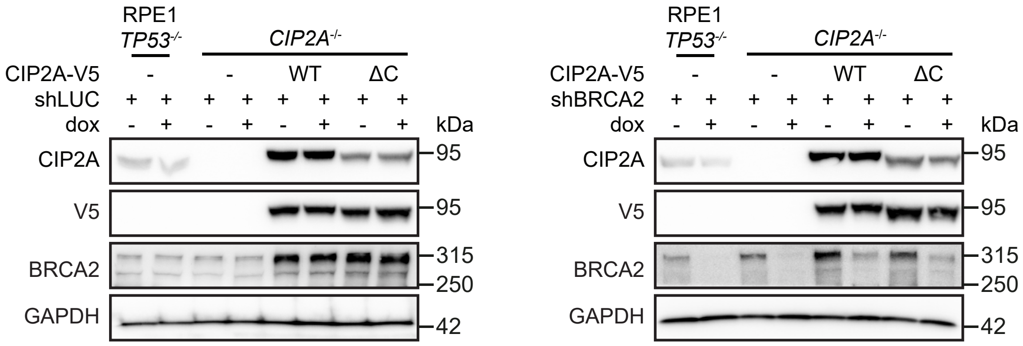


**B**

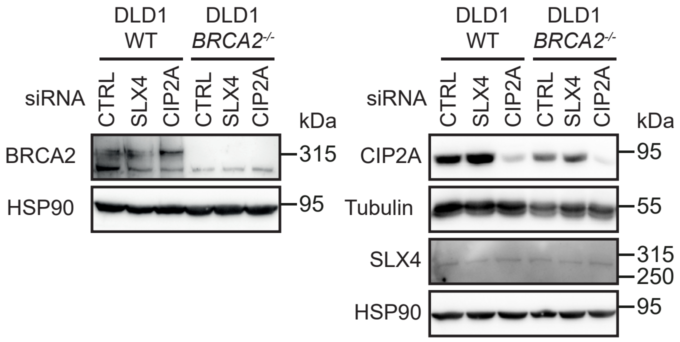
